# Multiplexed Targeted iprm-PASEF Assays for MALDI Mass Spectrometry Imaging of Lipids, NAD^+^, and Related Metabolites in CD38 Knockout Mice

**DOI:** 10.1101/2025.05.25.655991

**Authors:** Charles A. Schurman, Joanna Bons, Prasanna Vadhana Ashok Kumaar, Sriya Cheruvu, Jingqi Fang, Andrea Roberts, Genesis Vega Hormazabal, Rebeccah R. Riley, Nannan Tao, Eric Verdin, Birgit Schilling

**Author notes:** Corresponding Author: Birgit Schilling, Ph.D. Professor, Buck Institute for Research on Aging Director, Mass Spectrometry and Proteomics Core Buck Institute for Research on Aging Novato, CA 94945.

## Abstract

Mass spectrometry imaging (MSI) maps the molecular landscape of tissues; however, confident *in situ* identification and accurate quantification of analytes by matrix-assisted laser desorption/ionization (MALDI)-MSI remain challenging. We present a new targeted imaging methodology, imaging parallel reaction monitoring parallel accumulation-serial fragmentation (iprm-PASEF), which utilizes trapped ion mobility spectrometry (TIMS) and MALDI for assessment of metabolites and lipids on the fragment ion level in tissue. We investigated liver tissues from wild-type and CD38 knockout (CD38^-/-^) mice, as CD38 contributes to age-related nicotinamide adenine dinucleotide (NAD^+^) decline and direct detection of NAD^+^ within tissue remains challenging. The iprm-PASEF workflow enabled confident differentiation of lipid isomers, resolving phosphatidylinositols at *m/z* 857.52 and confirming the presence of PI 36:4(16:0/20:4) and absence of PI 36:4(14:0/22:4) at the fragment ion level. We confirmed a more than 2-fold increase in NAD^+^ in CD38^-/-^ liver tissues of male and female animals (n = 3 per genotype and sex) and decreased adenosine diphosphate ribose (ADPR) through relative quantification of characteristic fragment ions, confirming alteration of NAD metabolism in these animals. This approach provided confident and specific MS2-based identification and relative quantification of fragment ions in MSI experiments, with consistent fold changes across multiple characteristic fragment ions and biological replicates.

## Introduction

Mass spectrometry imaging (MSI) is a rapidly advancing application of mass spectrometry (MS) allowing for the unbiased spatial assessment of a variety of analytes and biological samples. MSI promises the capability to image thousands of molecules, such as metabolites, lipids, peptides, proteins, and glycans in tissues, while preserving their original spatial tissue localization.[1] This allows for the combination of the power, sensitivity, and specificity of mass spectrometry with spatial molecular details that can reveal novel biological regulation. Access to spatially resolved molecular features (proteins, glycans, lipids and metabolites) is instrumental to understand high tissue heterogeneity with localized hot spots of tissue damage, specifically during aging, (age-related) diseases and clinical applications. Using global proteomics/glycomics/lipidomics or metabolomics these spatial features are very challenging to capture, and indeed, often remain undetected or non-significant in global (‘bulk-style’) analyses. On the other hand, a variety of MS imaging approaches exist with multiple sources and ionization strategies, such as desorption electrospray ionization (DESI), laser ablation electrospray ionization (LAESI), liquid extraction surface analysis (LESA), liquid-micro junction-surface sampling probe (LMJ-SSP), infrared matrix-assisted laser desorption electrospray ionization (IR-MALDESI), and nanoscale secondary ion mass spectrometry (NanoSIMS).[2, 3] Another popular approach for MSI is matrix-assisted laser desorption/ionization (MALDI)-MSI which utilizes a deposited matrix-enhanced analyte ionization in place with laser ablation. MALDI-MSI allows for the discrete collection of pixel-specific mass spectra that are then assembled to generate a visual representation of the molecular species throughout the imaged region.

Several of the existing imaging techniques are limited to MS1 level analysis and do not utilize precursor ion fragmentation to increase confidence in analyte identification or improve quantification specificity and accuracy using fragment ion (MS2) level data. In these cases, analyte identification may rely on the concept of “accurate mass” assignment, developed for Electrospray Ionization High-performance Liquid Chromatography Mass Spectrometry (ESI-HPLC-MS/MS) approaches for instruments with high mass accuracy such as Fourier Transform Mass Spectrometry (FTMS) or Fourier Transform Ion Cyclotron Resonance (FTICR) instrumentation.[4] For imaging approaches utilizing MALDI-Time of Flight (TOF) - MSI, mass accuracy alone may not be sufficient to claim “accurate mass” without utilizing additional separation modalities such as trapped ion mobility spectrometry (TIMS) to provide an additional dimension of separation that aid in identification of target species. However, TIMS separation can be susceptible to spurious changes in instrument calibration and environmental effects that may slightly alter ion mobility and change the assigned analyte identification based on time or geographic location, or be complicated by isomeric and isobaric species that share molecular weight or ion mobilities.[5, 6] Powerful applications of MSI, such as DESI-MS, use a charged solvent spray to desorb and ionize analytes directly from the tissue surface through an ESI-like mechanism, which allows the tandem MS (MS2) fragmentation strategies traditionally used in HPLC-ESI-MS/MS to be applied in a spatially resolved manner.[7] However, DESI-MS can be affected by humidity and variable extraction efficiency, and its spatial resolution is limited by the continuous solvent flow and the size of the liquid bridge formed between the tissue sample and the DESI probe.[8] In addition, because DESI relies on *in situ* solvent extraction, it shares with HPLC-ESI-MS/MS a dependence on extraction conditions, a known source of variability for low-abundance, labile small molecules such as NAD^+^, but without chromatographic separation of co-extracted species prior to fragmentation.[9]

Since MALDI-MSI occurs through a direct ionization into the gas phase via laser ablation and ionization, and does not utilize liquid phases for possible separation, recent advances of MALDI-MSI utilize TIMS for ion separation based on an ion’s individual inverse reduced ion mobility (1/K_0_) values [10]. The addition of ion mobility separation to MALDI-MSI provides an orthogonal dimension of analyte separation, providing a reduction of complexity and specificity to MSI experiments differentiating isomeric lipids/metabolites with identical m/z but different 1/K_0_. Further, newer workflows, such as spatial ion mobility-scheduled exhaustive fragmentation (SIMSEF) [11], utilize TIMS to complete MS/MS but are limited in speed through reliance on data-dependent acquisition for MS/MS data collection. New targeted imaging parallel reaction monitoring (iprm) combined with parallel accumulation-serial fragmentation (PASEF) [12–15], use parallel reaction monitoring (PRM) for MS2 acquisition in a novel workflow termed iprm-PASEF. PRM is a mass spectrometry technique used for the quantification and identification of targeted analytes in complex biological samples that is valued for its high sensitivity, specificity, and accuracy in proteomic studies [16, 17]. For high-resolution PRM, precursor ions (at specific m/z and retention times) are isolated in a targeted way and fragmented. The resulting fragment ions are subsequently analyzed with high-resolution MS detectors, such as Orbitraps or time-of-flight detectors (Q-TOF)[17, 18]. This allows for multiplexed simultaneous analysis of all fragment ions with high resolution and high accuracy (post-acquisition), in contrast to low-resolution single reaction monitoring (SRM) or multiple reaction monitoring (MRM) which typically require extensive optimization of Q1/Q3 transitions [19, 20]. Due to the high resolution and post-acquisition refinement of fragment ion selection for quantification, PRM-MS generally is not as prone to interferences, and it is particularly efficient for fast developments of multiplexed assays. Here, we demonstrate the applicability of the recently introduced iprm-PASEF workflow for *in situ* analysis and quantification of lipids and metabolites (specifically NAD^+^) in animal tissues. This approach enables the acquisition of multiple targeted MS/MS spectra in a single imaging tissue MSI workflow, facilitating the development of multiplexed and targeted MS2-based spatial strategies.

NAD^+^ (nicotinamide adenine dinucleotide) metabolism is of key relevance for maintaining cellular health, playing crucial roles in energy metabolism, DNA repair, transcriptional regulation, immune function and inflammation.[21, 22] NAD^+^ itself is a central member of the large NAD metabolite family including the reduced form of NAD^+^, nicotinamide adenine dinucleotide NADH, nicotinamide adenine dinucleotide monophosphate (NADP^+^), nicotinamide adenine dinucleotide monophosphate reduced form (NADPH), nicotinamide mononucleotide (NMN), nicotinamide mononucleotide, reduced form (NMNH), nicotinamide riboside (NR), nicotinamide riboside reduced form (NRH), and nicotinamide (NAM), among others. Many clinical conditions implicate dysregulated NAD metabolism, including neurodegeneration, cancer, immune disorders, and cardio and metabolic diseases.[23–25] Aging and age-related diseases and cellular conditions, such as cellular senescence, are also associated with declines in NAD^+^ and disruptions in NAD^+^-related metabolic processes, positioning NAD^+^ as a significant target for biomedical research.[26, 27] In the liver, reduced NAD^+^ levels have been associated with non-alcoholic fatty liver disease (NAFLD) and other age-related metabolic dysfunctions.[28, 29] CD38, is a potent enzymatic consumer of NAD^+^, and shows increased expression with age in several tissues.[21, 26, 30–33] In liver tissue, CD38 is predominantly expressed in Kupffer cells, the resident macrophages of the liver sinusoids, where its activity drives cellular senescence and inflammation with aging.[27, 29, 34] Thus, CD38 is considered as a major driver of the age-related NAD^+^ decline.[35] Notably, CD38 knockout mice (CD38^-/-^) exhibit higher mitochondrial NAD^+^ levels in the liver and resistance to age-related NAD^+^ decline compared to aged, wild-type (WT) controls, suggesting improved metabolic regulation and protection against aging-associated metabolic deterioration.[30]

Additionally, NAD^+^ supplementation has emerged as a promising therapeutic strategy to enhance cellular function and mitigate age-related decline.[36, 37] Given the importance of hepatic NAD metabolism in systemic metabolic regulation, understanding how NAD⁺ and its related metabolites are spatially distributed within the liver could have significant therapeutic implications. Quantification of NAD^+^ is a technical challenge given its extreme lability and the extensive sample processing required for extraction from tissues [38, 39] leaving a need for more straightforward assays with less handling. Additionally, there is specific interest in the spatial distribution of this metabolite, and several non-MS methodologies have been used to image NAD^+^ in tissue. Genetically encoded fluorescent sensors can report NAD^+^ levels or NAD^+^/NADH ratios with cellular resolution but require transgenic expression or transfection and are therefore restricted to engineered model systems, whereas phosphorus-31 magnetic resonance spectroscopic imaging (^31^P-MRSI) can map NAD content non-invasively in humans but requires ultra-high-field instrumentation and provides spatial resolution far too coarse to resolve tissue microstructure [39, 40]. MS imaging can in principle overcome both limitations by detecting endogenous NAD^+^ directly in tissue sections at micrometer resolution. However, NAD^+^ and its metabolites are structurally related and labile, and can generate overlapping ions or in-source fragments (e.g., ADPR-derived species), which complicates confident assignment when MSI relies on MS1 data alone. Thus, we employed our newly optimized iprm-PASEF assays to spatially map and relatively quantify lipids, NAD^+^, and related metabolites in liver tissues from wild-type (WT) and CD38 knockout (CD38^-/-^) mice with minimal sample processing, isolating NAD^+^ from related species by both ion mobility and *m/z* prior to confirmation of its identity through characteristic fragment ions. Here, we present our multiplexed iprm-PASEF methodology and demonstrate its capability for MS imaging, identification, and MS2-based relative quantification of lipids and metabolites directly within fresh-frozen liver tissue sections.

## Materials and Methods

### Chemicals and Samples

Formic acid, acetonitrile (ACN), trifluoroacetic acid (TFA), methanol, water (all HPLC-MS grade), 2,5-dihydroxyacetophenone (DHAP), 2,5-dihydroxybenzoic acid (DHB) N-naphthylethylenediamine dihydrochloride (NEDC), and chemical standards for nicotinamide adenine dinucleotide (NAD^+^), ADP-ribose (ADPR) and cyclic ADP-ribose (cADPR) were purchased from Sigma-Aldrich (St. Louis, MO, USA). Livers from mice with global deletion of CD38 (CD38^-/-^) (Jackson Laboratory, 129P2-Cd38tm1Lnd/J) and control, litter mate wild-type (WT) mice (18 months old) were collected at the Buck Institute for Research on Aging.[30] All animal studies were done in compliance with the IACUC at the Buck Institute for Research on Aging under protocol # A10285.

### Sample preparation for iprm-PASEF imaging

Livers were collected from male and female CD38^-/-^ mice and WT controls (n = 3 biological replicates each) and flash frozen with liquid nitrogen to preserve lipids and metabolites. Frozen livers were mounted directly to a cryotome (Leica, Germany) pre-chilled to -20 °C using a cellulose-based mounting media (M-1 Embedding Matrix, Epredia, Kalamazoo, MI). The portal vein was used to standardize the orientation of the left major lobe of each liver. Optical images of all analyzed sections are shown in Supp. Fig. 11. Tissue sections were cut at 10 µm and mounted on positively-charged Superfrost™ (Fisher) microscope slides; a minority of sections that could not be cut cleanly at 10 µm were cut at 12 µm. Section thickness was not recorded for individual sections. Tissue sections were desiccated and then stored under vacuum at −80 °C prior to matrix application. Chemical standards for NAD^+^, ADPR, and cADPR were dissolved in methanol and deposited onto bare microscope slides with a pipette at 1–10 μM for initial assessment of ion mobility and fragmentation (Fig. 4A, C) and at 200 μM for collision energy and laser shot optimization and for comparison of unlabeled and stable isotope-labeled NAD^+^ (**Supp. Figs. 7, 8, and 10**). Deposits were allowed to dry before matrix was applied by spraying. The HTX TM Sprayer M3 was used to apply matrix by aerosol [41]. For positive ionization mode experiments, 15 mg/mL DHAP or DHB in 80% ACN, 10 % methanol in water was applied with the following parameters: flow rate 120 μL/min, number of passes 6, temperature 70°C, nitrogen gas pressure 10 psi, pattern CC, velocity 1200 mm/min, drying time 5 s. For negative ionization mode experiments, 7 mg/mL NEDC in 70% methanol in water was applied with the following parameters: flow rate 150 μL/min, number of passes 10, temperature 70°C, nitrogen gas pressure 10 psi, pattern HH, velocity 700 mm/min, drying time 0 s.

### MALDI-MS Imaging Analysis

#### MALDI-TIMS-MS1 acquisitions

MALDI-MS Imaging was performed using a Trapped Ion-Mobility Time-of-Flight Mass Spectrometer, the timsTOF fleX MALDI-2 (Bruker Scientific, LLC, Bremen, Germany) equipped with a dual ESI/MALDI source. Survey MS1 scans for lipids and metabolites were acquired with single-scan spectra with the smartbeam 3D 10 kHz laser set to acquire 200 accumulated laser shots at 1 kHz. Images were acquired at a 20 μm pixel resolution for both polarity ion imaging experiments. Other laser parameters included 35% power scan range of 16 μm for X and Y, and resulting field size of 20 μm for X and Y. Additional instrument parameters for the studies included: an ion transfer time of 50 μs, a pre pulse storage time of 60-80 μs, a collision RF of 800-1000 Vpp, a collision energy of 10 eV, an ion energy in the quadrupole of 5.0 eV, a TIMS funnels 1 RF and 2 RF of 250 Vpp, and a multipole RF of 250 Vpp. Negative ionization mode acquisitions targeting lipids ranged from m/z 150-1100 and 1/K_0_ 0.4-1.8 Vs/cm^2^ with a 100-ms ramp time. Positive ionization mode acquisitions, tuned for small molecule metabolites specifically, ranged from m/z 100-1000 and 1/K_0_ 0.4-1.4 Vs/cm^2^ with a 200-ms ramp time. Prior to every acquisition, laser focus, including Z-depth, was calibrated and sample profile assessed within timsControl software using the Sample Carrier and Focus Adjustment tools, with automatic target-profile correction of laser height across the slide and tissue surface.[42] Additional calibration was performed for the m/z dimension with spotted red phosphorus via MALDI-MS and the mobility dimension with a known peptide mix (Agilent) via ESI-MS directly on the timsTOF fleX before each imaging acquisition.

#### iprm-PASEF acquisitions

iprm-PASEF was performed in both positive and negative ionization acquisition modes. Main instrument settings were not adjusted between MS1 and MS2 acquisitions. Both modes used a mobility-based isolation strategy that isolated precursor ions by a 1/K_0_ window first followed by isolation by m/z. Collision energy was determined by a linear fit based on precursor mobility ranging from 35 eV at 1/K_0_ 0.8 Vs/cm^2^ to 60 eV at 1/K_0_ 1.6 Vs/cm^2^. No isolated precursor ions had a mobility greater than 1/K_0_ 1.6 Vs/cm^2^. iprm-PASEF precursor isolation settings including m/z and mobility ranges can be found in **Supplemental Table 1.** The effect of CID energy on the target class and ion mobility value is further investigated in **Supp. Fig. 1**.

#### MALDI-MSI data analysis (MS1)

After MS acquisition, the data was imported into SCiLS Lab Pro v2025a or v2025b (Bruker Scientific, LLC, Bremen, Germany) for initial MS1 data visualization and iprm-PASEF target list assembly. MetaboScape v.2025, (Bruker, Germany) was used to annotate analytes using existing or customized libraries to build precursor ion list based on MS1 m/z, isotopic pattern, and ion mobility values (**Supplemental Table 2**). Subsequently, iprm-PASEF data was used to confirm analyte identification by comparing fragmentation spectra to LipidMaps and the MS-DIAL-TandemMassSpectralAtlas for both positively- and negatively-charged analytes as well as fragment spectra for synthetic metabolite standards collected via HPLC-ESI-MS/MS (see below). Data was analyzed using SCiLS Lab Pro v2025a and v2025b, and Data Analysis v4.0 software (Bruker, Germany). iprm-PASEF data were further used for MS2-level relative quantification of selected fragment ions. Regions of interest (ROIs) were selected independently for each tissue section based on the areas of highest MS1 signal for the target analyte (ADPR or NAD^+^) in the survey scans, and iprm-PASEF data were acquired within these ROIs. For each animal, the quantitative value was the mean fragment ion intensity of all non-zero pixels within the ROI; pixels without detected signal were excluded, as technical dropout cannot be distinguished from analyte below the limit of detection. Non-normalized intensities were used, as low-molecular-weight matrix peaks can distort TIC or RMS normalization in this *m/z* range.[43, 44] Per-animal means were compared between WT and CD38^-/-^ animals by two-sided Student’s t-test in Prism v10.4.2 (GraphPad), with n = 3 biological replicates per genotype and α = 0.05. Quantitative results are provided in Supplemental Tables 3–4.

### HPLC-ESI-MS/MS Analysis

In order to validate detection of endogenous labile analytes in iprm-PASEF, including NAD^+^, ADPR, and cADPR, direct HPLC-ESI-MS/MS analysis of pure standards, obtained from Sigma Aldrich (N0632 for NAD^+^, A0752 for ADPR, and C7344 for cADPR), was completed to generate reference fragmentation spectra to connect HPLC-ESI-MS/MS to MALDI-MSI-MS/MS analysis of the same analytes (**Supp. Fig. 2**). MALDI-MS/MS spectra of the same standards were also acquired directly by iprm-PASEF (**Fig. 4A; Supp. Figs**. **7–8**). HPLC-ESI-MS/MS was included because publicly available reference spectra for these metabolites (e.g., METLIN, MoNA, MS-DIAL) are predominantly ESI-derived, so ESI spectra of the same standards provide a direct link between library spectra and MALDI-generated fragment ions. Standards at 10-micromolar concentration were injected onto a Thermo Hypercarb column (5 µm, 100 × 2.1 mm) using a Vanquish UHPLC system. Chromatographic separation was carried out over a 12-minute gradient using solvent A (7.5 mM ammonium acetate with 0.05% ammonium hydroxide in water) and solvent B (0.05% ammonium hydroxide in acetonitrile) as follows: 0–1 min, 5% B; 1–6 min, linear gradient to 60% B; 6.1–7.5 min, 90% B; 7.6–12 min, re-equilibration at 5% B. Mass spectrometric detection was performed on a Thermo Q Exactive mass spectrometer operated in positive ion mode using a targeted SIM-PRM acquisition strategy. Targeted SIM (tSIM) scans were acquired across a mass range of m/z 50–750 at 70,000 resolution, AGC target of 1e6, and maximum injection time of 100 ms. PRM scans were performed at a resolution of 17,500, with an AGC target of 5e5, maximum injection time of 55 ms, and stepped normalized collision energies (NCE) of 20, 35, and 30. Data were visualized using Thermo Freestyle software.

## Results

Fresh livers were collected from male and female 18-month-old CD38^-/-^ and control (WT) mice (with n=3 for each genotype for each sex), flash frozen, sectioned onto charged glass slides and prepared for MS imaging with the Bruker timsTOF fleX MALDI-2 mass spectrometer. CD38^-/-^ animals were chosen for the spatial profiling of lipids and metabolites (**Fig. 1A**), due to their well-characterized increase in hepatic NAD^+^ levels with age compared to control (WT) counterparts[30]. Samples were first subjected to unbiased MS1 imaging in both negative and positive ionization modes for survey scans of the whole tissue cross-section. Preliminary identification of interesting candidate MS1 precursor ions, based on accurate m/z, isotope pattern, and ion mobility values (1/K_0_), was completed with the T-ReX^3^ feature finding algorithm in SCiLS Lab, and features were annotated directly in SCiLS Lab by connecting to the MetaboScape 2025 REST API feature (**Fig. 1B, Supplemental Table 2**). To prioritize, candidate precursor ions that showed a predictive score difference of >0.6 comparing CD38^-/-^ and WT liver tissues with the area under the receiver operating characteristic curve (AUROC) analysis were considered for the subsequent iprm-PASEF analysis. The iprm-PASEF target lists were assembled so that the candidate ions did not overlap in its 1/K_0_ range based on the MS1 survey scans. For the iprm-PASEF acquisitions each candidate precursor ion was isolated first by their ion mobility 1/K_0_ value in the TIMS cell, then by its m/z value in quadrupole Q1, and then finally fragmented in the Q2 collision cell. The MS/MS spectra were acquired in this targeted, sequential, and spatially resolved manner (**Fig. 1C**). All fragment ions for each analyte shared the same mobility isolation window as their precursor ion but had unique m/z values corresponding to lipid or metabolite fragment ions (as annotated in existing databases/spectral libraries, see methods). Spatial maps were generated for each fragment ion that matched the spatial distribution of the corresponding precursor ion.

**Figure 1.**
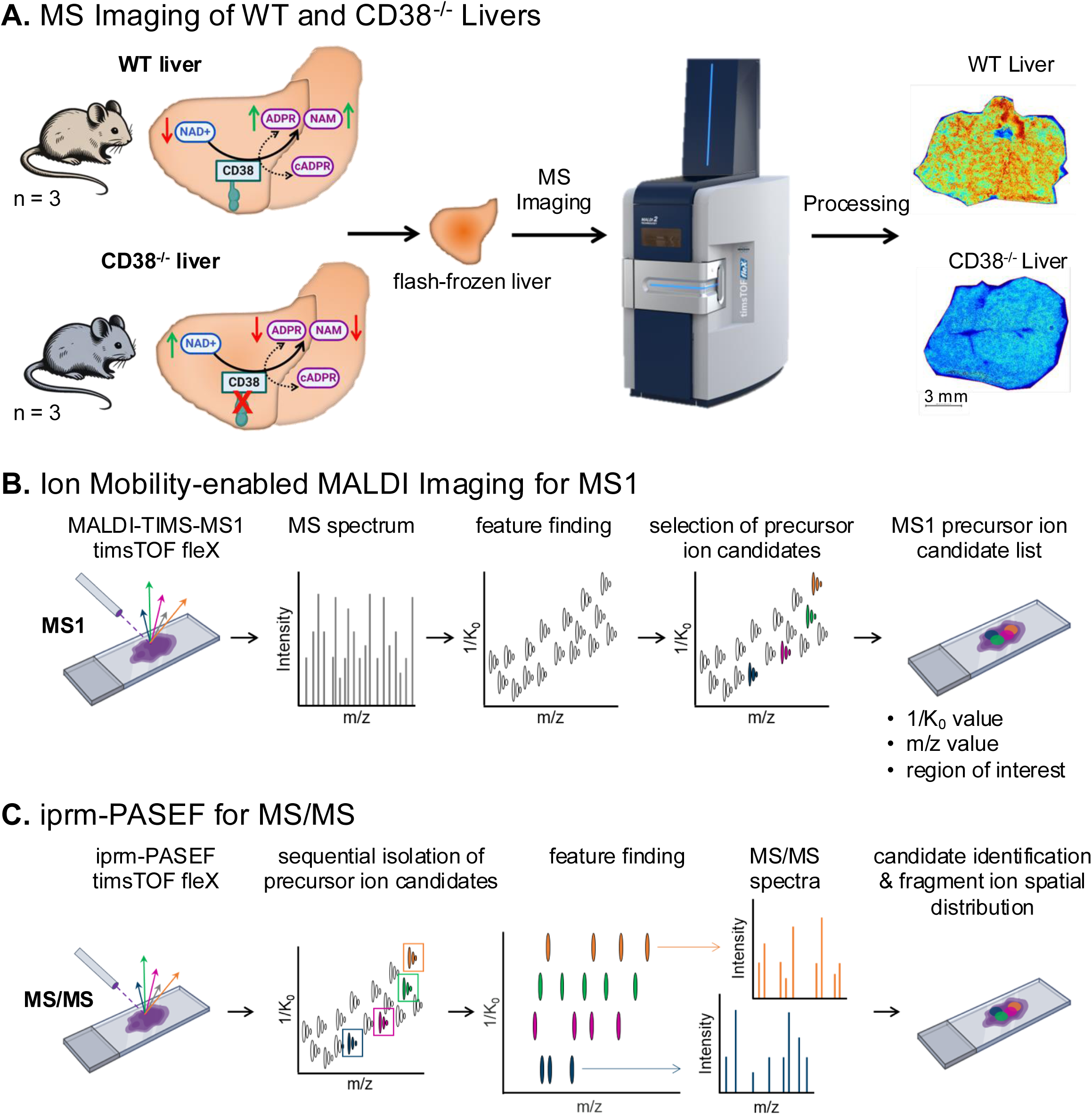
Experimental design for targeted MALDI-mass spectrometry imaging by iprm-PASEF of selected lipids and metabolites upon CD38 deletion in mouse liver. A. CD38 is a potent enzymatic consumer of NAD^+^ and CD38 knockout mice (CD38^-/-^) showed improved metabolism and lower cellular senescence in the liver, possibly through boosting the beneficial metabolic effects of NAD^+^. B. As a first discovery step, liver tissues from wild-type (WT) and CD38^-/-^ mice (n = 3 each) were subjected to collision cross section-enabled matrix-assisted laser desorption/ionization (MALDI)-mass spectrometry Imaging to spatially profile the lipidome and metabolome. Tissues sections were analyzed on a timsTOF fleX, MALDI-2 mass spectrometer by MALDI-trapped ion mobility spectrometry (TIMS)-MS in both negative and positive ionization modes. Survey imaging scans were processed in Bruker SCiLS Lab using the T-ReX^3^ feature finding algorithm and the MetaboScape software. A list of precursor ion candidates, including lipids and metabolites, was generated. C. As a second validation step, targeted iprm-PASEF assays were developed and optimized in SCiLS Lab and timsControl software, extracting information on 1/K_0_, m/z and region of interest from the MALDI-TIMS-MS1 scans. Each precursor ion was ion mobility-aided isolated and *in situ* fragmented sequentially. Collected iprm-PASEF data were analyzed in SCiLS Lab to apply the feature finding algorithm and generate MS/MS scans. This enabled to confirm lipid and metabolite candidate identification, to map the spatial distribution of their fragment ions in the tissues, and to perform multiplexed MS2-based quantification of the fragment ions.

To demonstrate the ability for isolation and accurate identification of individual lipids and metabolites from complex biological matrices by iprm-PASEF, in our data sets, we first targeted a well-characterized and annotated feature at m/z 885.55 corresponding to phosphatidylinositol PI(18:0/20:4) as reported in previous studies.[45] In negative ion mode this feature at m/z 885.55 was abundant in the MS1 spectra for all liver sections imaged (**Fig. 2A**). In the mobility domain, the species at m/z 885.55 in the MS1 survey scan matched a prominent feature at 1/K_0_ 1.47 Vs/cm^2^ (**Fig. 2B**). This feature was then isolated (**Supp. Fig. 3**) based on its unique 1/K_0_ (ion mobility) and m/z value combination allowing distinct analysis of this molecular feature separated from other features present in the liver tissue. The isolated candidate precursor ion was fragmented resulting in multiple abundant fragment ions. Importantly, the fragment ions all shared the same 1/K_0_ value as the isolated precursor ion, which allowed for generating a MS/MS spectrum of the fragmented [M-H]^-^ precursor ion accordingly (**Fig. 2C**). Fragment ions were annotated based on m/z values from existing databases and spectral libraries for lipid fragmentation. Major characteristic fragment ions corresponded to structures, such as stearic acid C(18:0) at m/z 283.26 and arachidonic acid C(20:4) at m/z 303.23 as well as several ions containing unique combinations of the phosphatidylinositol head group with the characteristic lipid tails at m/z 419.26, 581.31, and 599.32, among others (**Fig. 2C**). Combined, these confidently confirmed the identity of the precursor ion as PI(18:0/20:4). At the given MS/MS settings the precursor [M-H]^-^ ion at m/z 885.55 was still detectable even though at a significantly lower intensity than in the MS1 spectrum. To note, iprm-PASEF acquisition settings can be carefully fine-tuned to provide high quality MS/MS information ensuring confident ion detection, as illustrated for the collision energy applied for m/z 885.55 fragmentation (**Supp. Fig. 4**). MS/MS fragmentation was completed in a spatial raster manner at distinct tissue locations to generate the MS/MS imaging data set of fragment ions spatially across the liver sections (**Fig. 2D**). This showed that each fragment ion generated from one precursor ion displayed the same spatial distribution, sharing similar areas of high and low abundance across the tissue. Going forward, iprm-PASEF quantification can thus be performed on the fragment ion level for characteristic fragment ions increasing confidence in analyte assignment through precursor isolation and quantification based on characteristic fragment ions. This combination allows for more specific quantification by ignoring contributions from potential interfered or co-isolated species and also offers a more accurate measurement of analytes by ensuring measured abundance does not include contributions from interfered analytes that may be present on the MS1 level or have shared fragment ions on the MS2 level[11].

**Figure 2.**
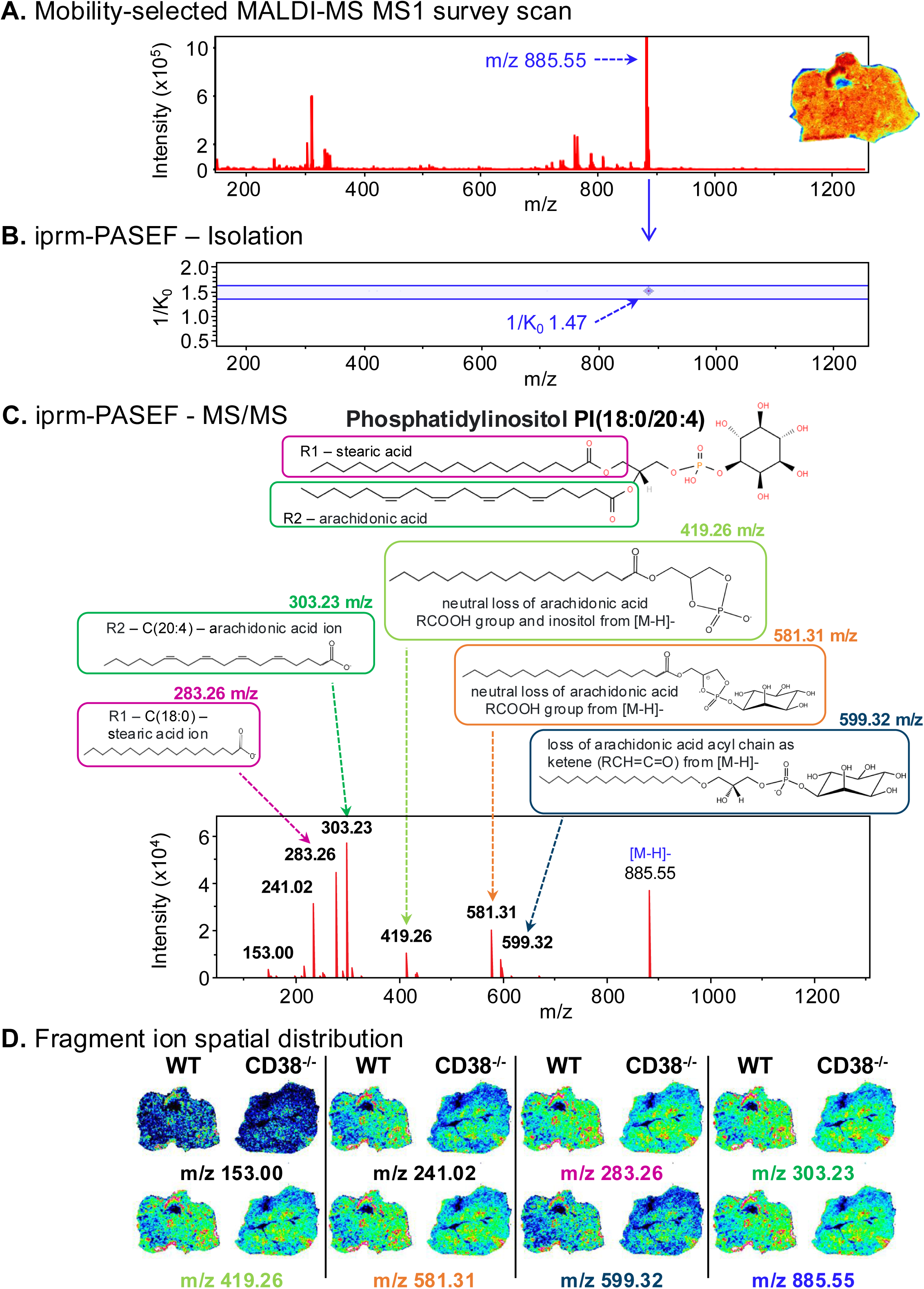
The iprm-PASEF strategy enables confident molecular identification with spatially resolved fragment ion distribution: example of Phosphatidylinositol PI(18:0/20:4). An ion mobility-selected survey MALDI-MS Imaging scan was acquired at the MS1 level to identify precursor ion candidates for targeted iprm-PASEF MS/MS-level analysis. Individual features of interest were selected based on their 1/K_0_ and m/z values. Up to 25 individual precursor ions, which were non-overlapping in 1/K_0_, could be simultaneously targeted for iprm-PASEF isolation and analysis. The iprm-PASEF MS/MS spectra for isolated and fragmented precursor ions were generated in a spatial manner in real-time. A. In CD38^-/-^ mouse liver, the composite survey MALDI-MS scan was acquired in negative ionization mode and showed the detection of a feature at m/z 885.55, that was highly abundant across the entire tissue section. B. This precursor ion candidate at m/z 885.55 and 1/K_0_ 1.47 Vs/cm^2^ was selected for targeted iprm-PASEF isolation and fragmentation. C. The generated MS/MS spectrum showed species-specific fragment ions used for both accurate molecular identification and quantification of Phosphatidylinositol PI(18:0/20:4). The identified fragment ions were spatially mapped for representative images of wild-type (WT) and CD38^-/-^ mouse liver tissue sections based on their m/z. The spatial distribution of the PI(18:0/20:4) fragment ions matched the distribution of the corresponding precursor ion.

The survey MS1 scans revealed the regions in the composite MALDI-MS spectrum where different lipid isotope patterns were observed. One area of interest was the m/z range from m/z 857 to m/z 861 (**Fig. 3A**). Initial annotation (based on databases) of the intense peak at m/z 857.52 returned two possible isomeric phosphatidylinositols PI(36:4), each with 36 carbon atoms and a total of 4 unsaturated double bonds, PI 36:4(16:0/20:4) and PI 36:4(14:0/22:4). However, an overlapping isotopic distribution was observed, as the third isotopic peak (m/z 859.53) showed higher intensity than one would expect for a PI 36:4 (PI 36:4), suggesting the presence of a PI 36:3 (at m/z 859.53). Moreover, the spatial pattern for this ion at m/z 859.53, as determined by MALDI-MSI analysis, differed from that at m/z 857.52 (data not shown). The iprm-PASEF analysis was completed to ascertain the identities of these related lipid species. First, the feature at m/z 857.52 was isolated and fragmented for analysis. Characteristic fragment ions for PI 36:4(16:0/20:4) were detected in the isolated mobility window at 1/K_0_ 1.4308 ± 0.0093 Vs/cm^2^ corresponding to palmitic acid C(16:0) at m/z 255.23 and arachidonic acid C(20:4) at m/z 303.23 (**Fig. 3B**). Many other fragment ions were also detected that further confirmed the identity of this lipid, including several head group fragment ions validating the identity of the phosphatidylinositol lipid family (**Supp. Fig. 5**). Some fragment ions detected in the iprm mobility window were unique to this MALDI-MSI iprm-PASEF experiment and did not match to annotated fragment ions from HPLC-ESI-MS/MS analysis of this lipid. Also, characteristic fragment ions of the potential isomer, PI 36:4(14:0/22:4), including features at m/z 227.20 (myristic Acid 14:0) and m/z 331.26 (adrenic acid 22:4), were not detected on the iprm mobilogram for this isolated feature (**Fig. 3B**). This enabled us to resolve its identity as PI 36:4(16:0/20:4) and confidently differentiate it from its potential isomer PI 36:4(14:0/22:4). Similar analysis was completed for the precursor ion at m/z 859.53 (**Supp. Fig. 6**) identifying this feature as a combination of two different PI isoforms (for PI 36:3, 3 double bonds), specifically: PI (18:1/18:2) and PI (16:0/20:3), which were distinct from the two Dalton lower mass identified at m/z 857.52 as PI 36:4(16:0/20:4) with 4 double bonds as described in Fig. 3 above. While iprm-PASEF has allowed for the confident identification of the composition of these complex lipid species, identifying the specific location of double bonds within the acyl tails generating positional isomers of each specific lipid is beyond the current capabilities of this application. Other methodologies such as oxygen or ozone induced dissociation or other chemical derivation techniques may in the future be employed within MALDI-MSI for these purposes.[46–48] These analyses of complex lipid species re-emphasize the importance of validation of analyte identities for accurate quantification utilizing fragmentation approaches that can delineate isobaric and isomeric species. Notably, overlapping isotopic envelopes of this kind cannot be resolved on the MS1 level alone, so that annotations returned by feature finding software such as MetaboScape remain preliminary. Because iprm-PASEF isolates each precursor ion by its ion mobility and m/z and then quantifies on fragment ions that are diagnostic for the individual species, specificity is retained even when the isolation window is not perfectly selective and more than one species is co-isolated.

**Figure 3.**
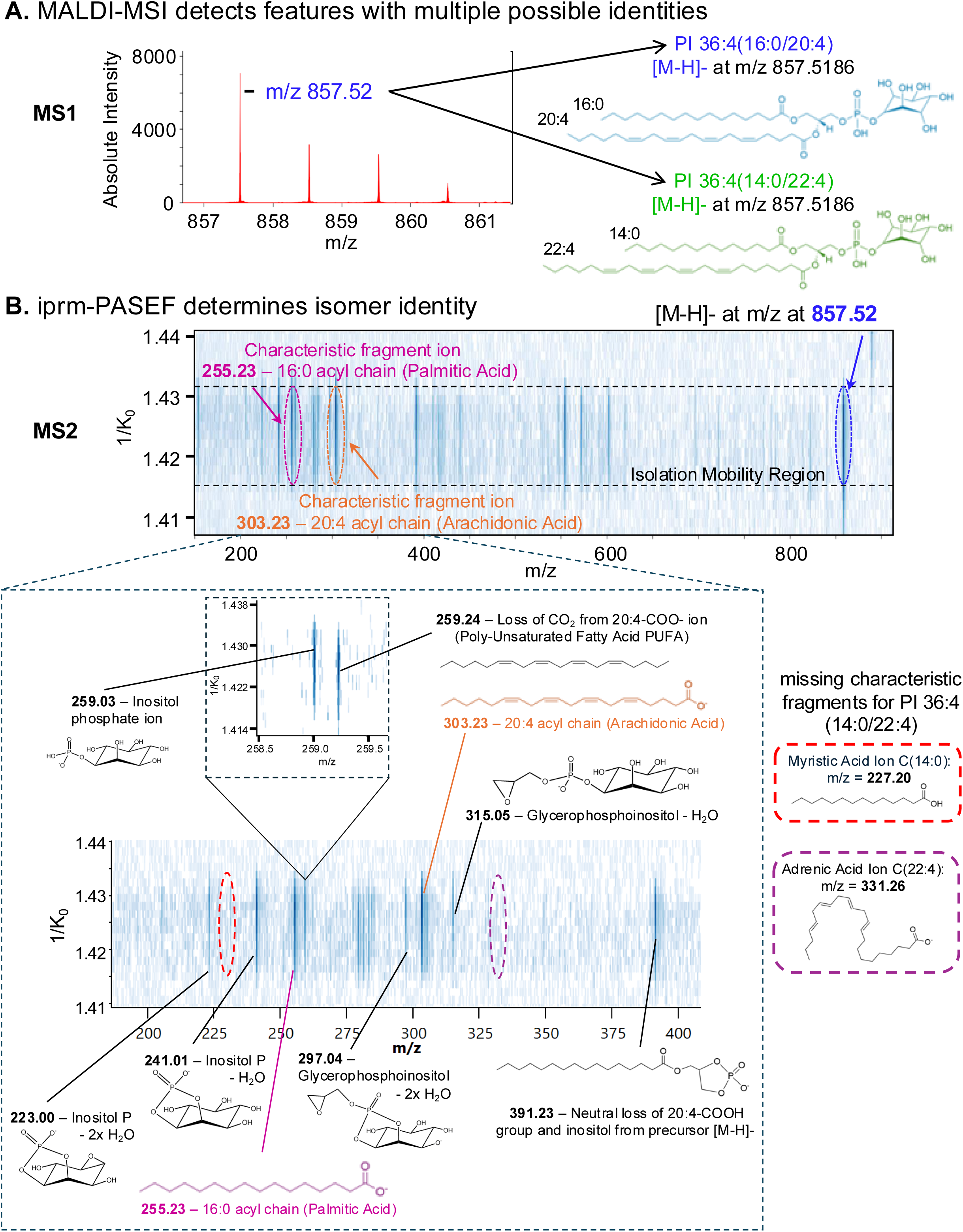
Detection of characteristic fragment ions for the spatially resolved identification of lipid isomers and isotopes in mouse liver tissues using iprm-PASEF. A. The zoom into the m/z range 857-861 of the composite survey MALDI-MS scan collected for CD38^-/-^ mouse liver tissue displayed an isotopic envelope for analytes (left). Potential lipid isomer discovery for the most intense ion at m/z 857.52 was initially annotated as PI 36:4, specifically as PI (14:0/22:4) and PI (16:0/20:4) by MetaboScape software. B. The iprm-PASEF fragmentation pattern of the precursor ion [M-H]^-^ at m/z 857.52 in the mobility isolation window 1/K_0_ 1.4308 ± 0.0093 Vs/cm^2^ showing the detection of two characteristic fragment ions, m/z 255.23 (pink) and m/z 303.23 (orange), as evidenced for the identification of PI 36:4 (16:0/20:4). No presence of characteristic fragment ions for PI 36:4 (14:0/22:4) confirmed the absence of this species.

CD38 is a multifunctional enzyme with a predominant role as an NAD^+^ glycohydrolase, catalyzing the hydrolysis of NAD^+^ to ADP-ribose (ADPR) and nicotinamide. Additionally, CD38 exhibits ADP-ribosyl cyclase activity, generating cyclic ADP-ribose (cADPR) from NAD^+^, though this constitutes a minor fraction of its total catalytic function. Furthermore, CD38 functions as a cADPR hydrolase, hydrolyzing cADPR to ADPR. Collectively, these enzymatic activities contribute to the regulation of NAD metabolism and calcium signaling, which are integral to cellular homeostasis. To identify optimal matrix conditions for MSI of NAD^+^ and its metabolites, we analyzed synthetic standards of NAD^+^, ADPR, and cADPR, individually spotted onto glass slides using various commonly used matrices. Dihydroxyacetophenone (DHAP) was used for positive ionization mode acquisitions and N-naphthylethylenediamine dihydrochloride (NEDC) was used for negative ionization mode. The matrix choice was not altered between MS1 and iprm-PASEF (MS2) acquisitions for the same analyte class on the same tissue or slide. A representative mobilogram overview of the cADPR, ADPR, and NAD^+^ standards highlighted their distinct ion mobility drift times (1/K_0_ values) and indicated efficient separation in the TIMS dimension, while *in situ* fragmentation of the standards generated MS/MS spectra for each metabolite using iprm-PASEF (**Fig. 4A**). The fragmentation patterns obtained through iprm-PASEF showed strong concordance with those derived from conventional HPLC-ESI-MS/MS (**Supp. Fig. 2**), where both methodologies detected characteristic fragment ions for each metabolite investigated.

**Figure 4.**
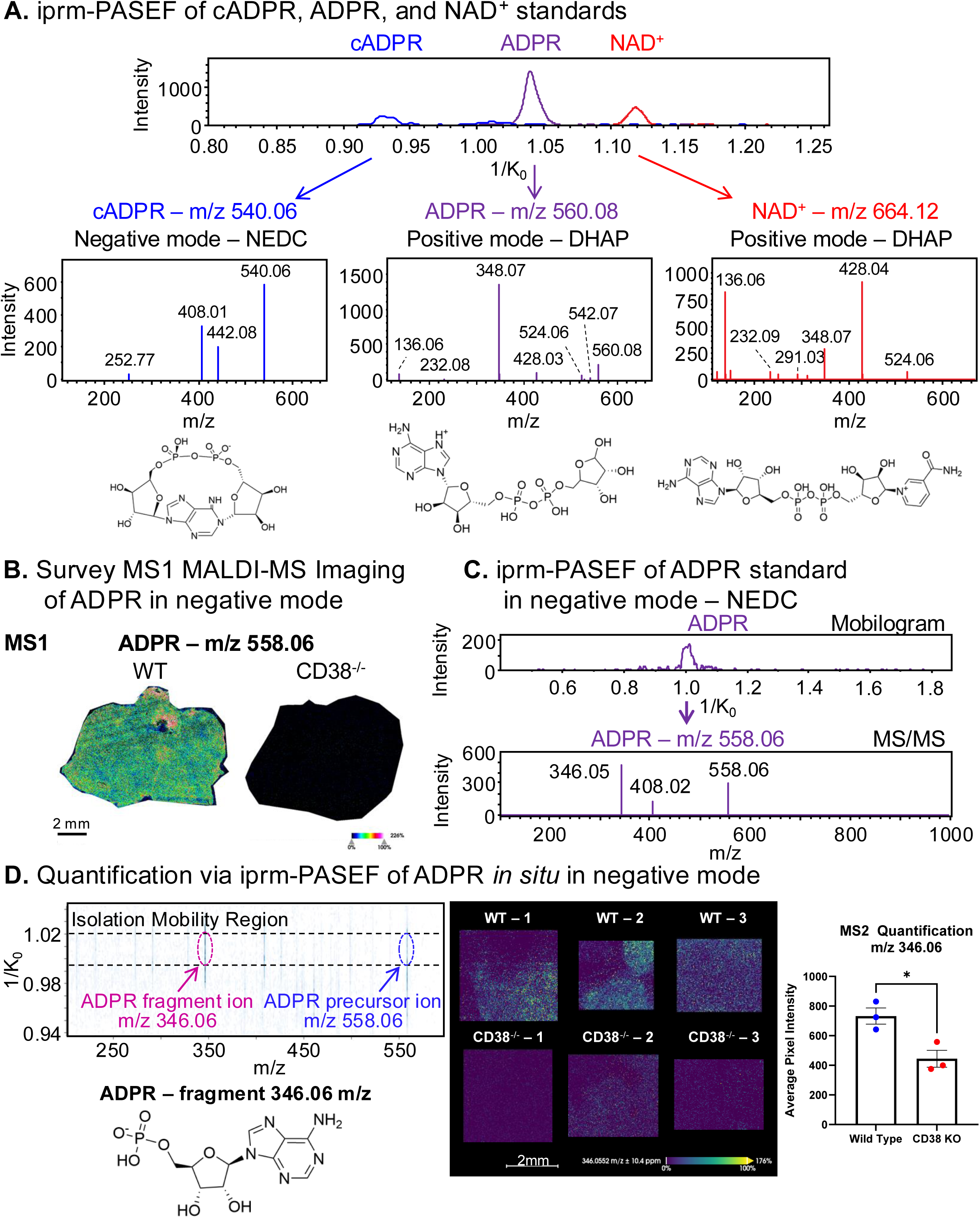
Confident identification of NAD^+^ and related metabolites in WT and CD38^-/-^ mouse livers by iprm-PASEF. A. The iprm-PASEF mobilogram and MS/MS spectra of synthetic cyclic adenosine 5′-diphosphate ribose (cADPR), ADPR, and nicotinamide adenine dinucleotide (NAD^+^) standards. Each standard was individually analyzed: in negative ionization mode (N-(1-naphthyl) ethylenediamine dihydrochloride (NEDC) matrix) for cADPR (1 μM) with precursor ion [M-H]^-^ at m/z 540.06, in positive ionization mode (2,5-dihydroxyacetophenone DHAP matrix) for ADPR (10 μM) with precursor ion [M+H]^+^ at m/z 560.08 and for NAD^+^ (10 μM) with precursor ion [M+H]^+^ at m/z 664.12. B. Spatial distribution of ADPR precursor ion (negative ionization mode, NEDC matrix, [M-H]^-^ at m/z 558.06) for representative WT and CD38^-/-^ mouse liver sections analyzed by MALDI-MS Imaging. C. The iprm-PASEF mobilogram and MS/MS spectrum of the synthetic ADPR standard analyzed in negative ionization mode (1 μM) with precursor ion [M-H]^-^ at m/z 558.06. D. ADPR was selected for targeted iprm-PASEF analysis in regions of interest selected from the areas of highest ADPR MS1 signal of WT and CD38^-/-^ mouse liver tissue sections. The fragmentation pattern of ADPR precursor ion [M-H]^-^ at m/z 558.06 in the mobility isolation window 1/K_0_ 1.0045– 1.0233 Vs/cm^2^ shows the detection of an ADPR fragment ion at m/z 346.06. This fragment ion was visualized *in situ* in the replicates of WT and CD38^-/-^ mouse liver and used for MS2-based relative quantification comparing the mean intensity of non-zero pixels within each ROI (ratio CD38^-/-^ vs. WT = 0.61, p = 0.022, corresponding to a 1.6-fold decrease; see Supplemental Table 3; * Student’s t-test p-value < 0.05 (n = 3 individual mice/biological replicates per condition).

Given that CD38 catalyzes NAD^+^ hydrolysis, we hypothesized that CD38^⁻/⁻^ liver would exhibit elevated NAD^+^ levels and a concurrent decrease in enzymatic by-products, such as ADPR and nicotinamide. To evaluate this, MALDI-MSI was performed for WT and CD38^⁻/⁻^ liver sections in negative ionization mode using NEDC matrix to monitor endogenous metabolites (n=3 biological replicates per genotype). The MS1 survey scans revealed a marked reduction in ADPR (m/z 558.06) in CD38^⁻/⁻^ liver, with almost negligible signal intensity in CD38^⁻/⁻^ liver compared to WT liver (**Fig. 4B**). To further validate this observation, an ADPR standard was analyzed again, now in negative ionization mode (**Fig. 4C**) which confirmed its characteristic fragmentation pattern. The optimized endogenous iprm-PASEF assay, incorporating MS2 quantification of the characteristic ADPR fragment ion at m/z 346.06 *in situ*, demonstrated a significant reduction in ADPR abundance in CD38 / tissue compared with WT (ratio 0.61, 1.6-fold decrease, p = 0.022; n = 3 biological replicates per genotype; Fig. 4D). These findings, consistent with previously reported results on ADPR, provided a proof-of-principle demonstration for CD38-mediated NAD metabolism in hepatic tissue and demonstrated the feasibility of the iprm-PASEF approach for *in situ* relative quantification of NAD^+^ metabolites at the MS2 fragment ion level.

Due to its highly polar and charged nature, NAD^+^ presents challenges with consistent ionization, which can impact detection sensitivity and quantification accuracy, particularly in the presence of ion suppression from lipids. In addition, NAD^+^ has in the past been shown to be prone to degradation thus this spatial imaging workflow may be advantageous due to very minimal sample handling[49]. In this study, we demonstrated efficient *in situ* detection and fragmentation of NAD^+^ in tissue using the iprm-PASEF approach. The iprm-PASEF workflow exhibited the capability to isolate and fragment the NAD^+^ precursor ion with [M+H]^+^ at m/z 664.12 in positive ionization mode with Dihydroxyacetophenone (DHAP) matrix (**Fig. 5A**). The presence of characteristic NAD^+^ fragment ions in the ion mobility isolation window, for the isolated precursor at m/z 664.12, including fragment ions at m/z 136.06, 232.08, 348.07, 428.04, and 524.06 confirmed the identity of the isolated metabolite as NAD^+^ and subsequently allowed for relative quantification of NAD^+^. For iprm-PASEF imaging of endogenous NAD^+^, acquisition parameters were selected from experiments with synthetic standards (**Supp. Figs. 7 and 8**). A collision energy of 35 eV maximized the abundance of the larger fragment ions at *m/z* 428.04 and 524.06 while limiting their further fragmentation into the smaller fragment ions at *m/z* 136.06 and 232.08, which are common to many adenine-containing metabolites (**Fig. 5B**). Fragment ion intensity increased markedly from 100 to 200 laser shots per pixel and continued to increase at higher shot numbers, but acquisition time scales directly with the number of shots (**Supp. Fig. 8F**). We therefore used 200 laser shots per pixel as a compromise between sensitivity and acquisition time, which was sufficient to resolve differences in NAD^+^ between genotypes. Two complementary correlation analyses were performed on the individual pixel intensities of these acquisitions. Across individual pixels, the five NAD^+^ fragment ions correlated strongly with one another, with a mean pairwise Pearson r of 0.85 at the optimized CE = 35 eV, consistent with their generation from a single isolated precursor ion (**Supplemental Table 6B**). In contrast, the correlation between the remaining precursor ion at m/z 664.12 and its fragment ions was high at low collision energy, where the precursor ion is only partially fragmented and the larger fragment ions at m/z 428.04 and m/z 524.06 dominate, and decreased with increasing collision energy as the precursor ion was progressively consumed and the larger fragment ions were themselves fragmented into the smaller product ions at m/z 136.06 and m/z 232.08 (**Supplemental Table 6C**).

**Figure 5.**
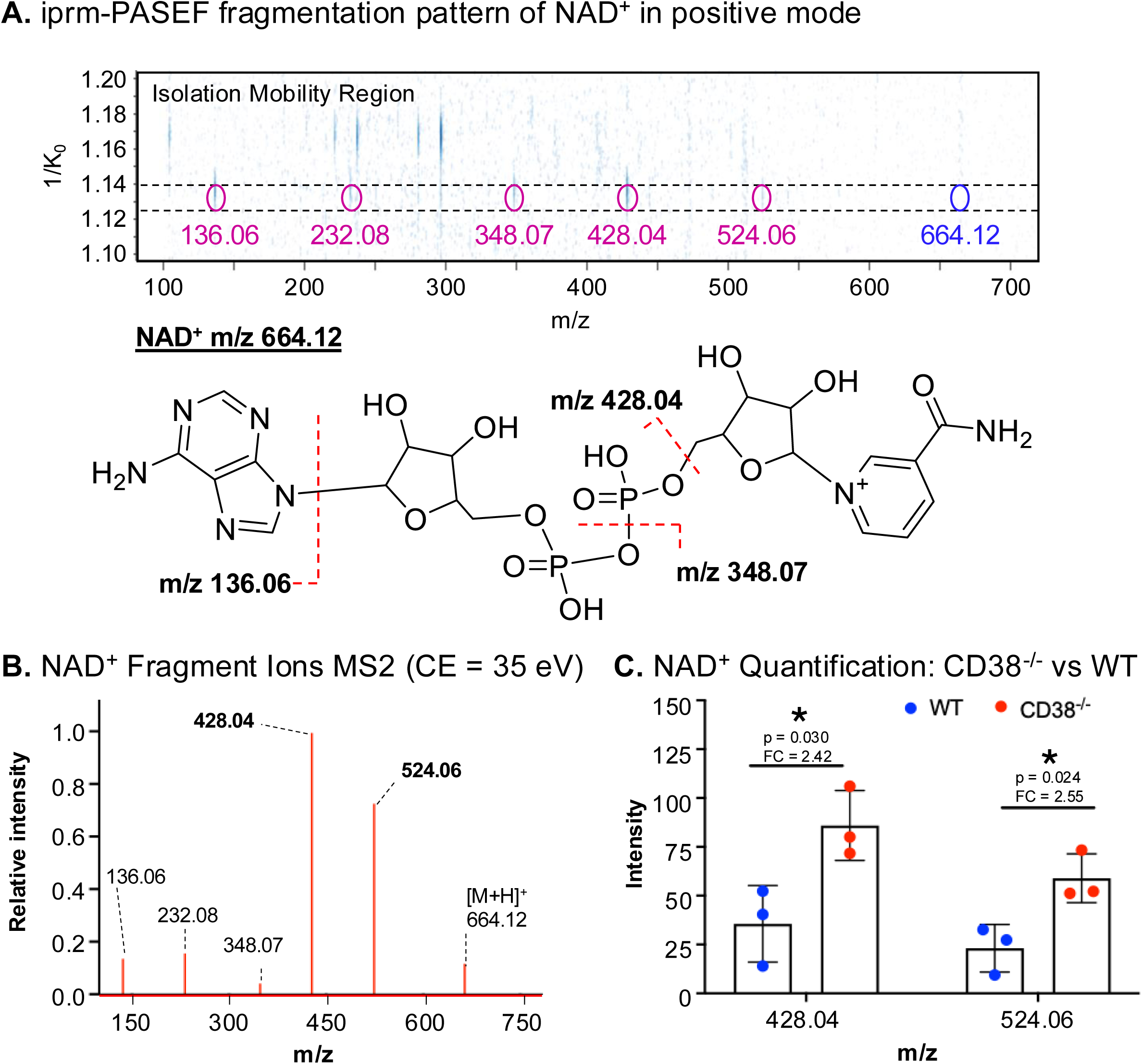
Quantification and *in situ* visualization of endogenous NAD^+^ fragment ions in WT and CD38^-/-^ mouse livers by iprm-PASEF. A. The iprm-PASEF fragmentation pattern of the endogenous NAD^+^ precursor ion [M+H]^+^ at m/z 664.12 in the mobility isolation window 1/K_0_ 1.1224-1.1400 Vs/cm^2^ showed the detection of fragment ions at m/z 136.06, m/z 232.08, m/z 348.07, m/z 428.04, and m/z 524.06. Cleavage sites giving rise to the fragment ions at *m/z* 136.06, 348.07, and 428.04 are mapped on the NAD^+^ structure. **B.** Using the acquisition parameters selected for tissue imaging (CE = 35 eV, 200 laser shots/pixel), the fragment ions at *m/z* 428.04 and *m/z* 524.06 were the most abundant and were selected for quantification. Because these fragments retain the ADP-ribose moiety and are also generated from ADPR (Fig. 4A), specificity for NAD^+^ is provided by isolation of the precursor ion by both *m/z* (664.12) and ion mobility (1/K_0_ 1.1224–1.1400 Vs/cm^2^), which excludes ADPR (*m/z* 560.08, 1/K_0_ ≈ 1.04 Vs/cm^2^). **C.** Relative quantification of NAD^+^ in liver sections from 3 female WT and 3 female CD38^-/-^ mice (Supp. Fig. 11). Per-animal means were compared by two-sided Student’s t-test (*m/z* 428.04: fold change 2.42, p = 0.030; *m/z* 524.06: fold change 2.55, p = 0.024; Supplemental Table 4). Bars show mean ± SD; points show individual animals. * Student’s t-test p-value < 0.05 (n = 3 individual mice/biological replicates per condition).

The MS2-level quantification and spatial imaging of each fragment ion revealed a significant increase in NAD^+^ levels in CD38^⁻/⁻^ liver tissues consistently in both males and females compared to WT, with around a 2.0-2.6 fold increase in liver tissue for each fragment ion (**Fig. 5C, Supp. Fig. 9**), which aligns with known NAD^+^ and CD38 biology. The consistency of elevation seen in multiple fragment ions used for relative quantification across multiple CD38^-/-^ livers increases confidence in quantification of this labile metabolite. These analyses are consistent with the established role of CD38 in NAD metabolism and support the use of iprm-PASEF for spatially resolved relative quantification.

## Discussion

The past decades have been marked by tremendous developments in omics technologies that have been instrumental for the in-depth exploration of complex biological systems, giving insights into molecular mechanisms and proposing novel clinical/therapeutic strategies. While very powerful, these technologies typically do not provide information about the native spatial context of biomolecules, which is highly relevant and critical to better appreciate the heterogeneity, architecture, and three-dimensional molecular organization of tissues. These needs have led to the rise of spatial omics technologies. Over the past five years, two spatial omics fields have been recognized as ‘Method of the Year’ by the journal *Nature Methods*, spatial transcriptomics in 2020[50] and spatial proteomics in 2024[51], demonstrating the powerful and crucial roles of these cutting-edge technologies to advance our understanding of biological complexity. Currently, even further advances are being made to improve the capabilities of these mass spectrometry imaging (MSI) techniques, especially in their ability to accurately identify and quantify complex biomolecules in place on tissue.

With respect to metabolomics and lipidomics, significant advances have been made in spatial mass spectrometry to investigate the spatial distribution of these molecules within tissues/organs. However, *in situ* identification remains very challenging. Here, we present a proof-of-principle study demonstrating the capabilities of imaging parallel reaction monitoring **–** Parallel Accumulation–Serial Fragmentation (iprm-PASEF) on lipids and metabolites, especially NAD^+^ and related metabolites, for high-resolution, spatially resolved targeted MALDI-MS/MS Imaging. MALDI-MSI with iprm-PASEF technology allows for high-confidence analyte identification and relative quantification in real-time on the MS2 fragment ion level during imaging experiments. This new workflow enables confident and accurate identification of complex and challenging molecules thanks to the acquisition of ion mobility-selected MS/MS spectra, while preserving their spatial information. Indeed, this strategy leverages the trapped ion mobility spectrometry (TIMS) technology to isolate molecular targets, first in the TIMS device based on their inverse reduced mobility (1/K_0_) and then in the quadrupole based on their mass-to-charge ratio (m/z). Isolated precursor ions are subsequently fragmented to collect high-resolution, full MS/MS spectra, while providing spatial information of both precursor and fragment ions. Ion mobility separation for lipid precursor ions is essential for proper identification and differentiation among many potential isoforms. MS2-level analysis further increases identity confidence through isolation and fragmentation of precursor ions of interest and subsequent detection of fragment ions that can be individually verified. The iprm-PASEF workflow utilizes the full capabilities of the timsTOF fleX and leverages the combination of ion mobility separation and precursor ion isolation and fragmentation to identify and quantify complex molecular features accurately. This workflow achieves label-free multiplexed and spatial multi-omic MS/MS imaging and quantification within a single acquisition.

The generated iprm-PASEF MS/MS spectra contain characteristic fragment ions that provide direct evidence for the *in situ* identification of lipids and metabolites present in tissues. The similar spatial distribution of the different fragment ions resulting from the same precursor ion provides an additional validation of the molecular identification. We showed that we can leverage the detection of characteristic fragment ions to decipher very complex and convoluted lipid profiles, as demonstrated for the well-studied and abundant Phosphatidylinositol PI(18:0/20:4) but also for more challenging and lower abundance features. The similarity in spatial patterns for identified fragment ions of PI(18:0/20:4) increased the confidence in isolated feature identification. Indeed, all fragment ions shared the same spatial pattern as the isolated precursor ion, signifying that all fragment ions were generated from a single precursor ion. Further, the ion mobility-resolved fragmentation pattern of the lipid PI 36:4(16:0/20:4) enabled highly confident identification of this feature and enabled its differentiation from its structural isomer PI 36:4(14:0/22:4). This exemplified the ability for iprm-PASEF to accurately resolve lipids utilizing complex fragmentation patterns for multiple features across a dynamic abundance range. The ability to accurately identify analytes in place during iprm-PASEF experiments removes the need for orthogonal mass spectrometry technologies to confirm analyte identifications, significantly increasing the ability of MALDI-MSI in discovery-based studies.

Besides confident identification, we demonstrated that the iprm-PASEF strategy can achieve quantification at the MS2 fragment ion level, using spatially resolved fragment ions, enabling more specific and sensitive performances. Similar to HPLC-ESI-MS/MS PRM assays developed for targeted (non-spatial) proteomic approaches, iprm-PASEF experiments can be refined post-acquisition to select the most intense, non-interfered, specific and selective, fragment ions for the analytes of interest for their specific quantification, while additionally providing a spatially defined context. This represents an asset in the context of complex and convoluted metabolome and lipidome analysis, as fragment ions differentiating co-fragmented species can be specifically used to ensure specific quantification. We developed and optimized new iprm-PASEF assays for *in situ* quantification of endogenous (and labile) NAD and ADPR metabolites in mouse livers. Through confident identification of these highly selective fragment ions, we demonstrated the ability for high confidence, specificity, and accuracy quantification for lipids and metabolites during MALDI-MSI experiments by utilizing the higher accuracy and identification confidence afforded by collecting data on the MS2 fragment ion level.

Our application of the iprm-PASEF workflow to CD38 and NAD^+^ biology demonstrates its power to better understand the underlying molecular mechanisms and address biological questions. NAD^+^ is a central redox coenzyme critical for cellular metabolism, supporting key oxidative and reductive reactions and serving as a substrate for NAD^+^-consuming enzymes such as sirtuins, PARPs, and the NADase CD38. With age, NAD^+^ levels decline in multiple tissues, including the liver and ovary, contributing to mitochondrial dysfunction and impaired metabolism[21, 26, 30–32]. Our findings recapitulate the role of CD38 as a major driver of this decline through its NADase activity, which hydrolyzes NAD^+^ primarily into nicotinamide and ADP-ribose (ADPR), as previously reported in independent studies [21, 26, 30–32, 52].

While global NAD^+^ quantification by HPLC-ESI-MS/MS is well established, understanding its spatial distribution is essential for elucidating compartment-specific metabolic regulation of NAD^+^ and related metabolites. Mass spectrometry imaging offers spatial resolution but, in some cases, faces challenges in detecting small, labile metabolites like NAD^+^ and ADPR due to ion suppression from abundant proteins and lipids, as well as difficulties in metabolite annotation. To overcome these limitations, prior studies have employed HRMS/MS and spiking of NAD^+^ standards onto control tissue[53, 54]. For instance, Lu et al. demonstrated that acidified methanol washes reduce ion suppression and, when combined with isotopic labeling, allow confident identification of NAD^+^ and related metabolites[55]. In this study, we utilized the iprm-PASEF strategy to map the spatial distribution of NAD^+^ and ADPR in wild-type and CD38 knockout mouse livers. MALDI-MSI of fresh-frozen tissue sections mounted on standard histology-grade charged glass slides allowed rapid assessment of endogenous metabolites with minimal sample handling and common reagents; indium tin oxide (ITO)-coated slides can be used instead, and their effect on ionization efficiency for these analytes warrants further investigation. Our approach using refined iprm-PASEF fragment ions enabled spatially resolved MS2-based quantification, *i.e.* specifically using the fragment ion at m/z 346.06 for ADPR (in negative ionization mode) and the abundant fragment ions using our optimized acquisition parameters, m/z 428.04 and 524.06 for NAD^+^ (in positive ionization mode). We were able to further conduct statistical analysis to confirm the significant increase of the NAD^+^ analyte and the significant decrease of the ADPR analyte upon CD38 deletion in mouse livers. Additionally, we optimized matrix concentration and application parameters for DHAP to enhance NAD^+^ ionization during MALDI MS Imaging experiments from both slide-spotted standards and directly from tissue sections, further improving MSI sensitivity and reliability. We also demonstrated that the collision energy determines the NAD^+^ fragmentation pattern and that the number of laser shots determines fragment ion intensity, which together informed the selection of acquisition parameters for the iprm-PASEF assays (**Supp. Fig. 7 and 8**). Further, we validated that for future studies where absolute quantification of metabolites is desired with iprm-PASEF, stable isotope-labeled (SIL) standards (**Supp. Fig. 10**) share the same isolation window for mobility isolation as non-heavy labeled standards or endogenous metabolites, ensuring that optimized instrument settings are equally applicable and should not overtly affect efforts for quantification. We note that the quantification reported here is relative quantification rather than absolute quantification, as no isotopically labelled internal standard was spiked into the tissue sections; comparisons were made for the same analyte across samples acquired under identical conditions. Additional validation including intra- and inter-day repeatability, linearity, and limits of detection for instrument performance, was beyond the scope of this proof-of-principle study and will be required before iprm-PASEF is applied to absolute quantification. Section thickness, which is known to influence MALDI ion yield [56], varied between 10 and 12 µm and was not recorded for individual sections. Although automatic target-profile correction compensates for height differences, an effect of section thickness on absolute ion intensities cannot be excluded; because the use of 12 µm sections was determined by sectioning quality rather than genotype, such variation would be expected to increase variance rather than bias the comparison between genotypes. The new iprm-PASEF workflow will allow for the study of more subtle and dynamic changes in metabolism, such as those occurring in female mice during the circadian cycle, aging, or metabolic stress. Such future applications will further expand the relevance of the iprm-PASEF approach to uncover spatial and temporal regulation of NAD⁺ metabolism under various physiological and pathological conditions.

## Conclusions

This work developed new targeted MS Imaging assays to measure and understand the spatial distribution of targeted analytes which is essential for elucidating compartment-specific metabolic regulation, here specifically of NAD^+^ and related metabolites. Overall, we envision the use of complementary, synergistic, and multi-modal approaches combining global HPLC-ESI-MS/MS and MALDI-MS/MS (iprm-PASEF) imaging to i) generate ESI-based spectral libraries to help interpret MALDI-MS/MS spectra and improve identification rate in MS-imaging assays, ii) inform on the spatial distribution of analytes of interest identified by HPLC-ESI-MS/MS experiments, iii) aid in integrating multi-omics measurements and data. This new iprm-PASEF workflow opens the way to new strategies for advanced spatial metabolomics, lipidomics, and proteomics – including *in situ* tryptic peptides[14] or any other proteolytic peptides/proteins, that will be pivotal in obtaining a more holistic and refined picture of the dynamic organization and regulation of biological systems. With the multiplexed capabilities of iprm-PASEF in performing high-quality identification and simultaneous quantification for multiple complex lipids and biologically relevant small molecule metabolites, we have demonstrated the importance and future opportunities of MS2-based spatial mass spectrometry imaging. Specific future interests could be in spatial mapping of tissues highlighting therapeutic drug distribution, regional biomarker responses upon interventions, and overall tissue heterogeneity during aging and age-related diseases.

## Data Availability

The spatial MALDI-MSI and liquid chromatography mass spectrometry raw MS data sets have been uploaded to the Center for Computational Mass Spectrometry, MassIVE, and can be downloaded using the following FTP link: ftp://MSV000097855@massive-ftp.ucsd.edu or via the MassIVE website: https://massive.ucsd.edu/ProteoSAFe/private-dataset.jsp?task=4dbd4a7e3a4d4f66b90399a693974e2f (MassIVE ID: MSV000097855).

[Note to the reviewers: To access the data repository MassIVE (UCSD) for MS data, please use: Username: MSV000097855_reviewer; Password: winter].

## Acknowledgements

We would like to acknowledge support from the following NIH funding sources: NIAMS R21 AR084303 (PI Schilling), NIA P01 AG066591 (PI Ellerby), NIA T32 AG000266 (PI Ellerby, fellowship to J. Fang). NIH Office of the Director S10 OD038264 (PI Schilling). We also acknowledge support from the Glenn Foundation for Medical Research (fellowship to C.A. Schurman). The authors would like to acknowledge Bruker for their technical support and generous access to instrumentation.

## Author Contributions

C.A.S and J.B. contributed to the study design, data acquisition and interpretation, manuscript and figure preparation. C.A.S led the revision and overall writing/editing of the manuscript. P.V.A.K. contributed to data acquisition and analysis, manuscript and figure preparation, and editing of the manuscript. S.C. assisted with sample acquisitions and analysis. Authors J.F., A.R, G.V.H., and R.R.R. contributed to the murine study design and sample acquisition. N.T. contributed to study design, provided resources and data acquisition and manuscript editing. E.V. contributed to study design. B.S. implemented the study design and conceptual development, provided resources, contributed to data interpretation and manuscript preparation and revision.

## Supplemental Material

**Supplemental Figure 1. Effect of CID energy depending on target ion mobility and chemical class.** The three metabolite and four lipid species investigated in this study are represented. Dots are color-coded depending on the chemical class: orange corresponds to lipids and purple to metabolites. The grey dotted line represents the linear regression used to determine the calculated collision energy (CE) based on the precursor mobility: 35 eV at 1/K_0_ 0.8 VS/cm^2^ to 60 eV at 1/K_0_ 1.6 VS/cm^2^. For the targeted analytes, ADPR, cADPR, and NAD^+^ metabolites were initially fragmented at lower collision energies (CE = 41.7 eV, 41.7 eV, and 45.4 eV, respectively) than the lipids PI(16:0/20:4), PI(18:1/18:2), PI(16:0/20:3), and PI(18:0/20:4) (CE = 54.7 eV, 54.8 eV, 54.8 eV, and 55.9 eV, respectively; see Supplemental Table 1). CE’s may be further optimized per analyte for preservation of different fragment ion intensity distributions.

**Supplemental Figure 2. HPLC-ESI-MS/MS spectra of NAD+, ADPR, and cADPR standards. A.** Analysis by HPLC-ESI-MS/MS of pure NAD+ synthetic standard on the Thermo Q Exactive Orbitrap mass spectrometer enabled the detection of several characteristic fragment ions of NAD+ that matched fragment m/z features detected by iprm-PASEF in MALDI-MSI for both the pure standard and the endogenous metabolite in fresh-frozen liver tissues **B.** ADPR and **C.** cADPR synthetic standard also led to the identification of characteristic fragment ions that matched those detected by iprm-PASEF.

**Supplemental Figure 3: iprm-PASEF isolation and fragmentation. A.** Features were first identified in the MS1 MALDI composite spectrum and spatial heatmaps were investigated. **B.** Mobility 1/K_0_ value for the feature of interest at m/z 885.55 was extracted from the mobilogram from the full untargeted MS1 acquisition. **C.** Using both 1/K_0_ and m/z, the precursor ion was isolated from surrounding molecular species. **D.** Fragmentation of the precursor ion generated fragment ions that shared the exact same ion mobility range.

**Supplemental Figure 4: Evaluation of the impact of the collision energy value on endogenous phosphatidylinositol PI(18:0/20:4) fragmentation pattern.** CD38^-/-^ mouse liver tissue was submitted to iprm-PASEF analysis in negative mode, targeting the precursor ion with [M-H]^-^ at m/z 885.55 and 1/K_0_ 1.47 Vs/cm^2^ that corresponds to phosphatidylinositol PI(18:0/20:4). Collision energy (CE) was ramped from 30 eV to 65 eV. A-F. MS/MS spectra of the endogenous phosphatidylinositol PI(18:0/20:4). G. Boxplots showing the distribution of the pixel intensities of the endogenous phosphatidylinositol PI(18:0/20:4) fragment ions at m/z 241.02, m/z 283.26, m/z 303.23, m/z 419.26, m/z 581.31, and m/z 599.32 as well as precursor ion at m/z 885.55 at the different CE tested (see **Supplemental Table 5**). Empty pixels were excluded. The black diamond represents the mean value. Points beyond 1.5 × IQR of the quartiles are shown individually and represent 0.3–4.0% (median 1.0%) of non-zero pixels per ion and collision energy. This revealed that CE = 30 eV was inefficient to fragment the precursor ion. From CE = 45 eV, fragment ions were detected, but the precursor ion was the most intense for CE ranging from 45 eV to 55 eV. Fragment ions at m/z 283.26 and m/z 303.23, that correspond to the stearic acid ion and the arachidonic ion respectively, were the most intense fragment ions.

**Supplemental Figure 5: Detected fragment ions from the precursor ion isolated at m/z 857.52 and 1/K_0_ 1.4308 ± 0.0093 Vs/cm^2^.** Annotation of the detected ions to known fragment ion m/z from the species PI 36:4(16:0/20:4) confirmed the lipid identity based on characteristic fragments for R-groups Palmitic Acid (m/z 255.23) and Arachidonic Acid (m/z 303.23). Multiple head group fragments, including a prominent ion at m/z 241.01 representing an inositol phosphate ion minus H_2_O, confirmed the lipid class identity. Pseudo MS/MS spectrum for the isolated species was assembled from detected fragment ions extracted from the iprm mobilogram.

**Supplemental Figure 6: Isomer resolution of co-fragmented species. A.** iprm-PASEF was used to isolate a precursor ion [M-H]- at m/z 859.53 in the mobility isolation window 1/K_0_ 1.4334 ± 0.0094 Vs/cm^2^. This species was preliminarily identified as PI 36:3(18:1/18:2) (green) from MS1 survey scans but also matched the precursor ion m/z for PI 36:3(16:0/20:3) (blue). **B.** The fragmentation pattern of the precursor ion [M-H]-at m/z 859.53 with a zoom on the m/z range 279.0-283.5 contained two detected characteristic fragment ions for R-groups Oleic Acid (18:1, m/z 281.25) and Linoleic Acid (18:2, m/z 279.23), while multiple head group fragments, including a prominent ion at m/z 241.01, representing an inositol phosphate ion minus H_2_O, confirmed the identification of the lipid species PI 36:3(18:1/18:2). Additional fragment ions at m/z 415.23 and 417.24 matched to fragments containing the PI headgroup and tails from the precursor [M-H]-PI 36:3(18:1/18:2). However, further investigation of additional fragment ions across a broader m/z range revealed strong signals for Palmitic Acid (16:0, m/z 255.23) and Dihomo-γ-linolenic Acid (DGLA) (20:3, m/z 305.23). This, along with other fragments containing acyl tails from PI 36:3(16:0/20:3), revealed that the precursor ion [M-H]-PI 36:3(16:0/20:3) was co-isolated with the precursor ion [M-H]-PI 36:3(18:1/18:2) and co-fragmented. Specific and accurate quantification can still be achieved by using characteristic fragment ions for each individual lipid.

**Supplemental Figure 7:** Evaluation of the impact of the collision energy value on NAD^+^ standard fragmentation pattern. Synthetic NAD^+^ standard was spotted on glass slides (200 µM) and DHAP matrix was sprayed for iprm-PASEF analyses in positive mode with precursor ion [M+H]^+^ at m/z 664.12 and 1/K_0_ 1.12 Vs/cm^2^. Collision energy (CE) was ramped from 20 eV to 55 eV. A-H. MS/MS spectra of NAD^+^ standard. I. Boxplots showing the distribution of the pixel intensities of NAD^+^ fragment ions at m/z 136.06, m/z 232.08, m/z 348.07, m/z 428.04, and m/z 524.06 as well as precursor ion at m/z 664.12 at the different CE tested (see Supplemental Table 6). Empty pixels were excluded. The black diamond represents the mean value. Points beyond 1.5 × IQR of the quartiles are shown individually and represent 0–6.3% (median 0.7%) of non-zero pixels per ion and collision energy. Standards were deposited by pipette and dried before matrix was sprayed. This revealed that CE = 20 eV was inefficient to fully fragment the precursor ion: it remained the most abundant ion, while CE ≥ 35 eV was enough to do so. At CE = 25-30 eV, both the precursor ion and the fragment ions were detected. In addition, the fragmentation pattern changed when increasing the CE: m/z 524.06 was the most abundant ion at CE = 25 eV, while it shifted to m/z 428.04 at CE = 30-45 eV and m/z 136.06 for CE ≥ 50 eV.

**Supplemental Figure 8: Evaluation of the impact of the number of laser shots on NAD^+^ standard fragmentation pattern.** Synthetic NAD^+^ standard was spotted on glass slides (200 µM) and DHAP matrix was sprayed for iprm-PASEF analyses in positive mode with precursor ion [M+H]^+^ at m/z 664.12 and 1/K_0_ at 1.12 Vs/cm^2^. The number of laser shots was ramped from 50 to 2000 at 1 kHz. **A-E.** The boxplots show the distribution of the pixel intensities of NAD^+^ fragment ions at m/z 136.06, m/z 232.08, m/z 348.07, m/z 428.04, and m/z 524.06 for the different laser shot numbers applied (see Supplemental Table 7). Empty pixels were excluded. The black diamond represents the mean value. Points beyond 1.5 × IQR of the quartiles are shown individually and represent 0–13.7% (median 4.7%) of non-zero pixels per ion at ≥200 laser shots; at 50 and 100 shots, too few pixels contained signal (9–33 per ion) for outlier percentages to be meaningful. Standards were deposited by pipette and dried before matrix was sprayed. At 50 and 100 laser shots, NAD^+^ fragments were barely detected. Above 200 shots, the intensity of the fragment ions increased upon enhancing the number of laser shots. **F.** Impact of the number of laser shots on the acquisition speed at 1 kHz laser repetition rate. 200 laser shots per pixel were used for tissue imaging as a compromise between fragment ion intensity and acquisition time.

**Supplemental Figure 9: Spatial Distribution and Quantification of endogenous NAD^+^ in livers of male CD38^-/-^ and WT mice via iprm-PASEF.** Spatial distribution of the four NAD^+^ fragment ions detected by iprm-PASEF in one WT and one male CD38^-/-^ mouse liver replicate show homogeneous distribution of NAD^+^ when detected in CD38^-/-^ tissue sections and low abundant signal in WT animals. B. Quantification of NAD^+^ abundance in WT and CD38^-/-^ liver sections in 3 male animals per genotype shows significant increases of fragments m/z 428.04 and m/z 524.06, corresponding to the endogenous NAD^+^ precursor ion [M+H]^+^ at m/z 664.12 & 1/K_0_ 1.1224-1.1400 Vs/cm^2^ in situ during imaging experiments with a fold change of ∼2x compared to WT (see Supplemental Table 4). For each animal, the non-normalized intensity of each fragment ion was averaged across all non-zero pixels within an ROI selected from the area of highest NAD^+^ MS1 signal and per-animal means were compared by two-sided Student’s t-test. Student’s t-test p-value < 0.05 (n = 3 individual mice/biological replicates per condition).

**Supplemental Figure 10: Light and heavy stable isotope-labeled NAD**^+^ **standards have the same ion mobility (1/K_0_) value and fragmentation pattern.** Synthetic light (**A,C**) and heavy (**B,D**) stable isotope-labeled NAD^+^ standards were spotted on glass slides (200 µM) and DHAP matrix was sprayed for survey MS1 and iprm-PASEF analyses in positive mode with light NAD^+^ precursor ion [M+H]^+^ at m/z 664.12 and heavy NAD^+^ precursor ion [M+H]^+^ at m/z 669.13. **A-B.** MS1 survey scans are shown for the region m/z 651 – 675 that contains both precursor ions of interest. This shows that the heavy-labeled ^13^C_5_-NAD+ standard is not contaminated by the light form, confirming that the fragment ions observed by iprm-PASEF analysis specifically results from the fragmentation of the heavy-labelled form. **C-D**. MS/MS mobility-m/z ratio heatmaps and spectra are displayed. Both standards presented similar 1/K_0_ values of 1.11-1.12 Vs/cm^2^ as well as similar fragmentation patterns with confident detection of diagnostic ions, *i.e.* m/z 232.08 and m/z 524.06 for the light form as well as m/z 237.10 and m/z 529.07 for the heavy form.

**Supplemental Figure 11: Optical Images of Liver Tissues. A)** Slide scan images of fresh frozen liver tissues used for iPRM-PASEF imaging captured at 2400 DPI for control (WT) and CD38 knock out (CD38^-/-^) mice from male and female. Livers were mounted using carboxymethylcellulose (CMC) and sectioned in the transverse plane through the left lobe attempting to preserve the portal vein structure (lower right) when possible. Scale bar = 10 mm. B) Representative image of serial liver sections mounted on the a standard 25 x 75 mm charged glass microscopy slide.

**Supplemental Table 1:** iprm isolation settings for targeted analytes

**Supplemental Table 2:** MetaboScape Lipid and Metabolite Annotations from MS1 Overview Scans of Mouse Liver Tissues

**Supplemental Table 3:** Quantification of ADPR fragment ions

**Supplemental Table 4:** Quantification of NAD^+^ fragment ions

**Supplemental Table 5:** Quantification of PI(18:0/20:4) precursor and fragment ions at different collision energies

**Supplemental Table 6:** Quantification of NAD^+^ precursor and fragment ions at different collision energies

**Supplemental Table 7:** Quantification of NAD^+^ fragment ions with different laser shots

