## Supplementary material for "Multiplexed Targeted iprm-PASEF Assays for MALDI Mass Spectrometry Imaging of Lipids, NAD^+^, and Related Metabolites in CD38 Knockout Mice": Supple Figure 1

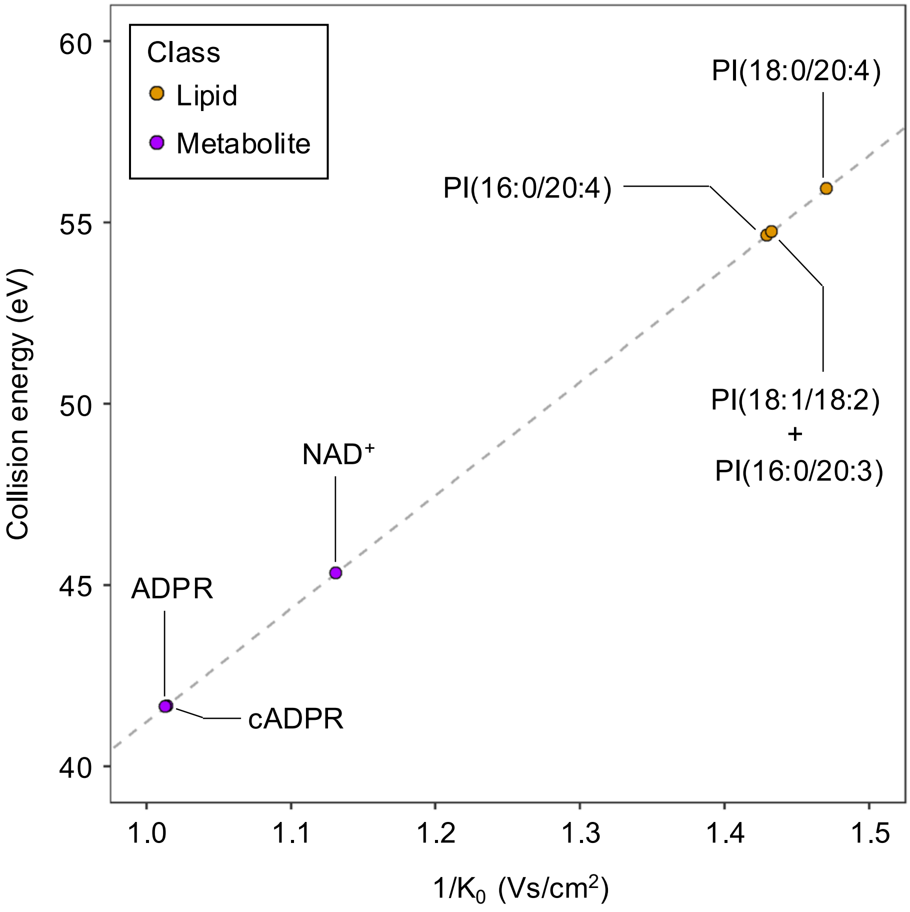


**Supplemental Figure 1: Effect of CID energy depending on target ion mobility and chemical class.** The three metabolite and four lipid species investigated in this study are represented. Dots are color-coded depending on the chemical class: orange corresponds to lipids and purple to metabolites. The grey dotted line represents the linear regression used to determine the calculated collision energy (CE) based on the precursor mobility: 35 eV at $1/K_0$ 0.8 $VS/cm^2$ to 60 eV at $1/K_0$ 1.6 $VS/cm^2$. For the targeted analytes, ADPR, cADPR, and $NAD^+$ metabolites were initially fragmented at lower collision energies (CE = 41.7 eV, 41.7 eV, and 45.4 eV, respectively) than the lipids PI(16:0/20:4), PI(18:1/18:2), PI(16:0/20:3), and PI(18:0/20:4) (CE = 54.7 eV, 54.8 eV, 54.8 eV, and 55.9 eV, respectively; see **Supplemental Table 1**). CE's may be further optimized per analyte for preservation of different fragment ion intensity distributions.
