## Supplementary material for "Multiplexed Targeted iprm-PASEF Assays for MALDI Mass Spectrometry Imaging of Lipids, NAD^+^, and Related Metabolites in CD38 Knockout Mice": Supple Figure 2

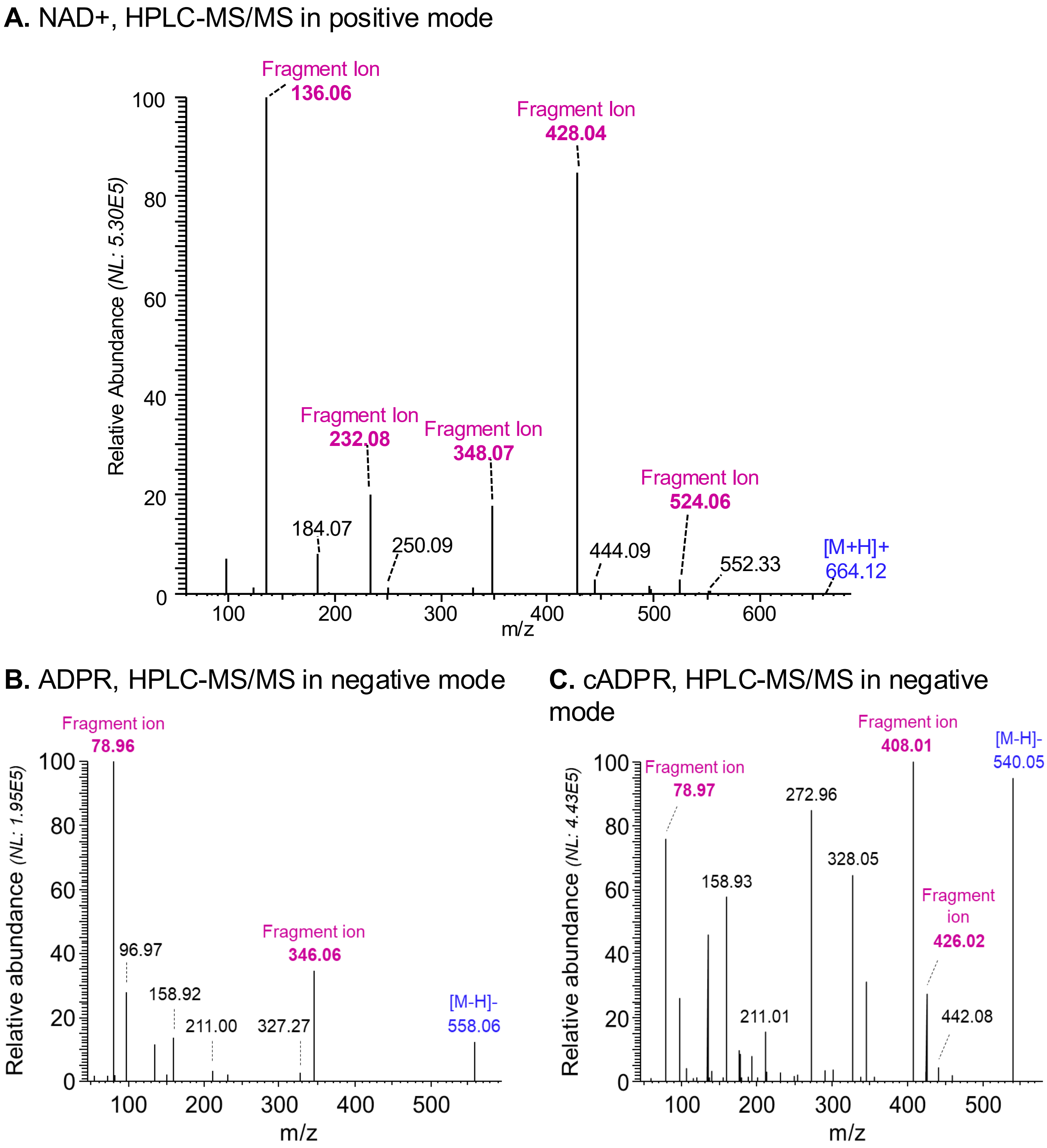


**Supplemental Figure 2: HPLC-ESI-MS/MS spectra of NAD+, ADPR, and cADPR standards. A.** Analysis by HPLC-ESI-MS/MS of pure NAD+ synthetic standard on the Thermo Q Exactive Orbitrap mass spectrometer enabled the detection of several characteristic fragment ions of NAD+ that matched fragment m/z features detected by iprm-PASEF in MALDI-MSI for both the pure standard and the endogenous metabolite in fresh-frozen liver tissues **B.** ADPR and **C.** cADPR synthetic standard also led to the identification of characteristic fragment ions that matched those detected by iprm-PASEF.
