## Supplementary material for "Multiplexed Targeted iprm-PASEF Assays for MALDI Mass Spectrometry Imaging of Lipids, NAD^+^, and Related Metabolites in CD38 Knockout Mice": Supple Figure 3

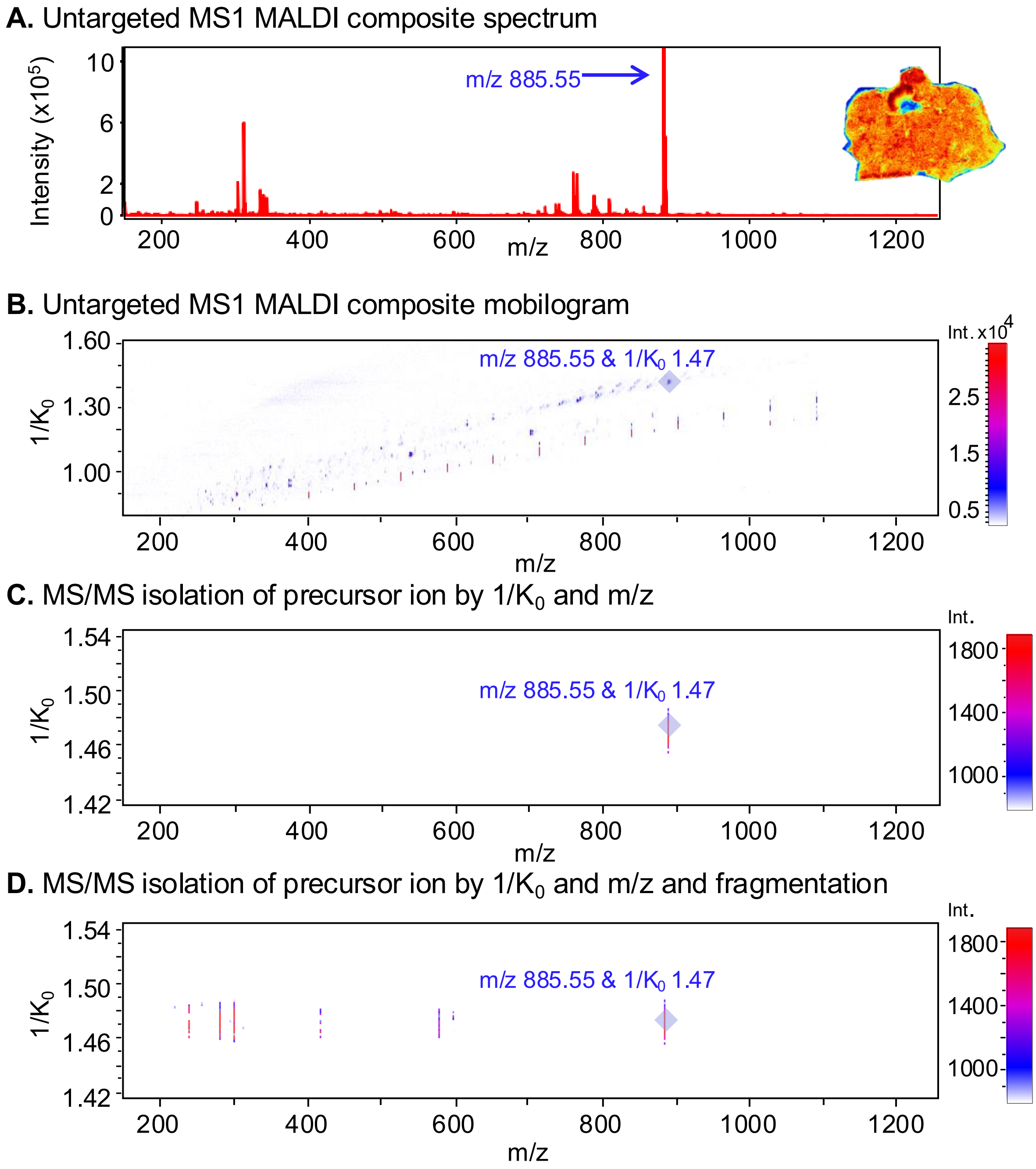


**Supplemental Figure 3: iprm-PASEF isolation and fragmentation. A.** Features were first identified in the MS1 MALDI composite spectrum and spatial heatmaps were investigated. **B.** Mobility $1/K_0$ value for the feature of interest at m/z 885.55 was extracted from the mobilogram from the full untargeted MS1 acquisition. **C.** Using both $1/K_0$ and m/z, the precursor ion was isolated from surrounding molecular species. **D.** Fragmentation of the precursor ion generated fragment ions that shared the exact same ion mobility range.
