## Supplementary material for "Multiplexed Targeted iprm-PASEF Assays for MALDI Mass Spectrometry Imaging of Lipids, NAD^+^, and Related Metabolites in CD38 Knockout Mice": Supple Figure 4

### Supplemental Figure 4

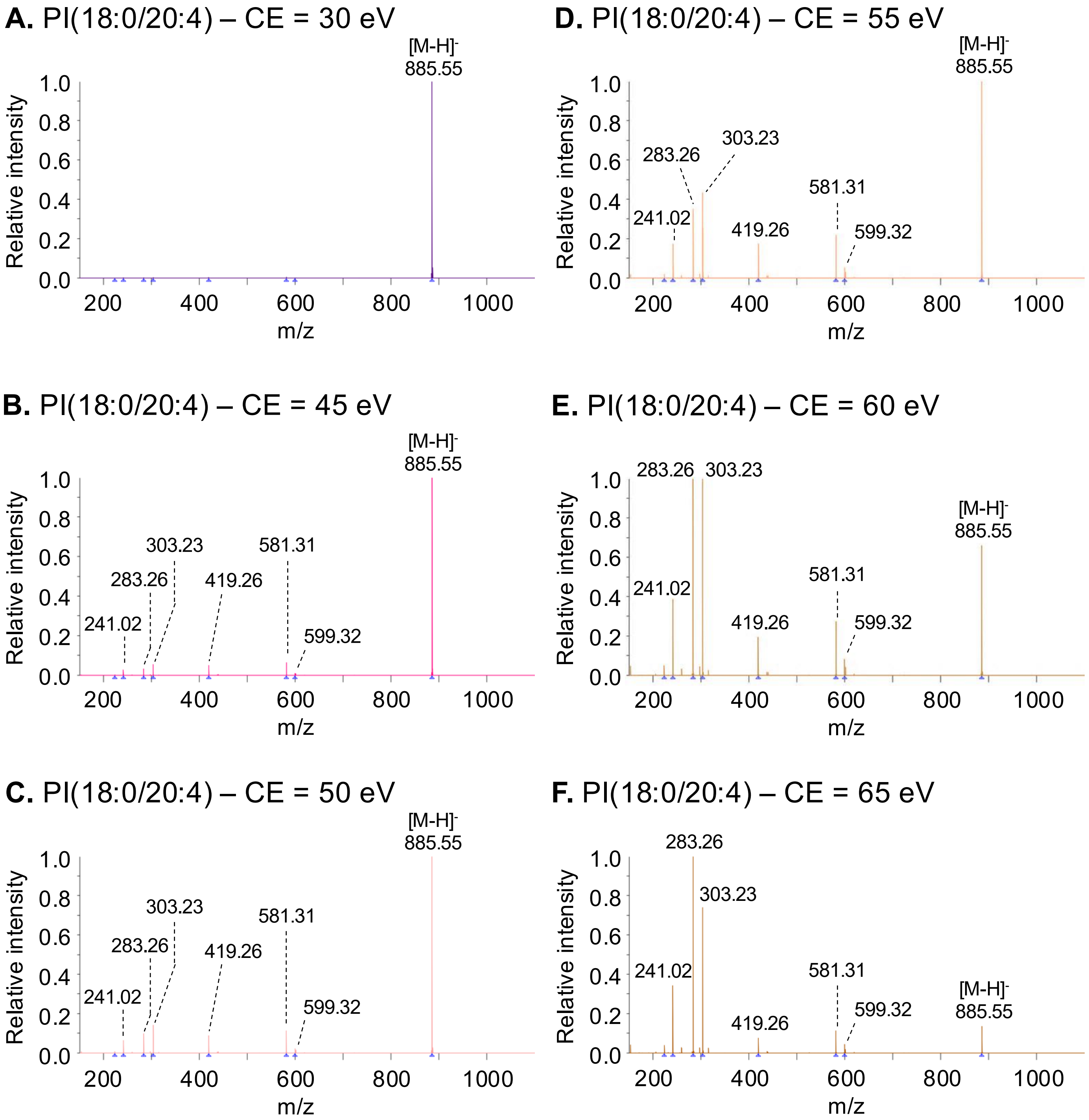

### Supplemental Figure 4

**G.** Precursor and fragment ion quantification

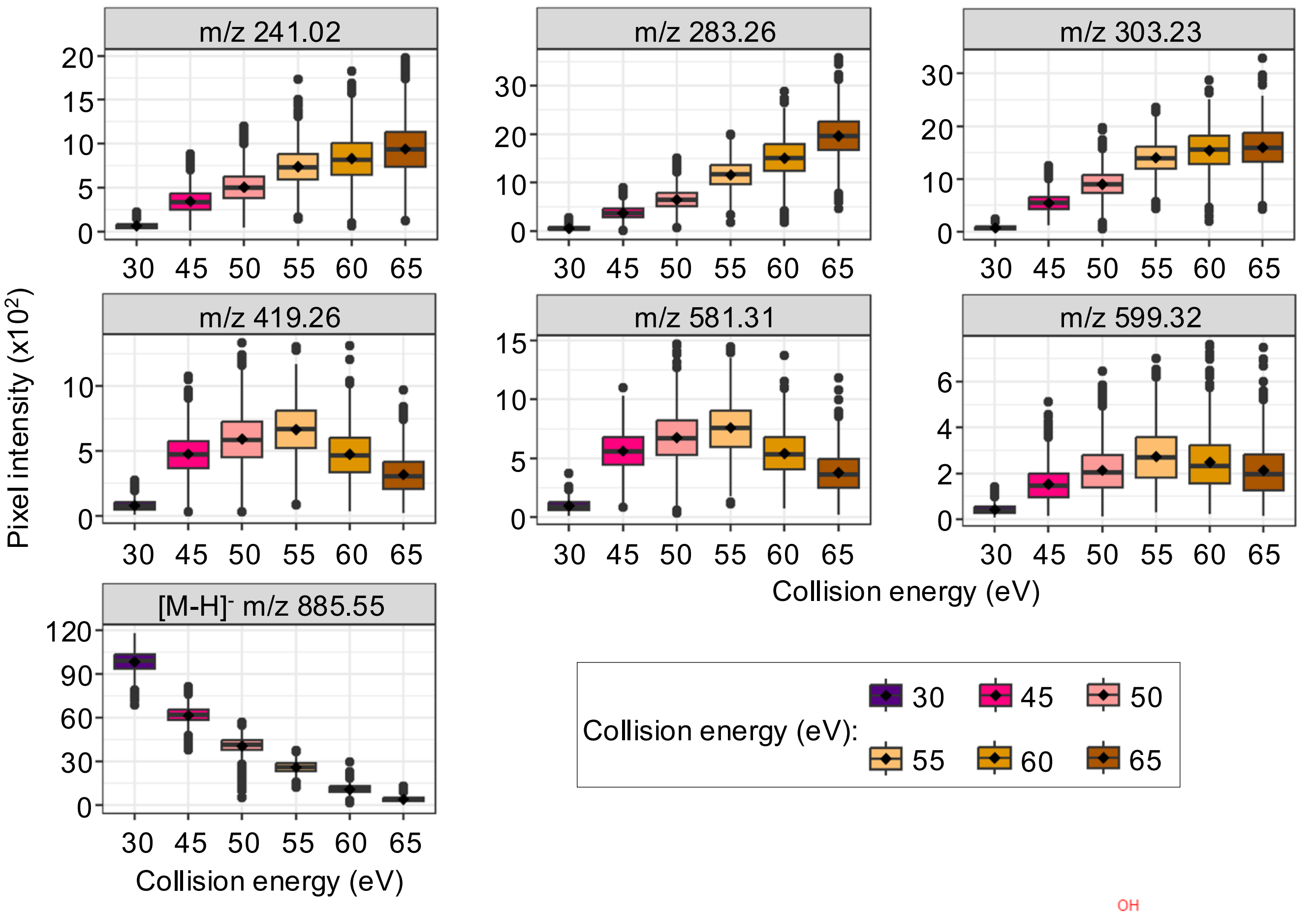


**Phosphatidylinositol PI(18:0/20:4)**

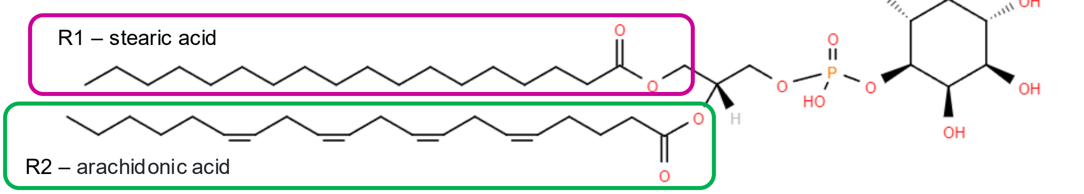


**Supplemental Figure 4: Evaluation of the impact of the collision energy value on endogenous phosphatidylinositol PI(18:0/20:4) fragmentation pattern.** CD38$^{-/-}$ mouse liver tissue was submitted to iprm-PASEF analysis in negative mode, targeting the precursor ion with $[M-H]^-$ at m/z 885.55 and $1/K_0$ 1.47 $Vs/cm^2$ that corresponds to phosphatidylinositol PI(18:0/20:4). Collision energy (CE) was ramped from 30 eV to 65 eV. A-F. MS/MS spectra of the endogenous phosphatidylinositol PI(18:0/20:4). G. Boxplots showing the distribution of the pixel intensities of the endogenous phosphatidylinositol PI(18:0/20:4) fragment ions at m/z 241.02, m/z 283.26, m/z 303.23, m/z 419.26, m/z 581.31, and m/z 599.32 as well as precursor ion at m/z 885.55 at the different CE tested (see **Supplemental Table 5**). Empty pixels were excluded. The black diamond represents the mean value. Points beyond 1.5 × IQR of the quartiles are shown individually and represent 0.3–4.0% (median 1.0%) of non-zero pixels per ion and collision energy. This revealed that CE = 30 eV was inefficient to fragment the precursor ion. From CE = 45 eV, fragment ions were detected, but the precursor ion was the most intense for CE ranging from 45 eV to 55 eV. Fragment ions at m/z 283.26 and m/z 303.23, that correspond to the stearic acid ion and the arachidonic ion respectively, were the most intense fragment ions.
