## Supplementary material for "Multiplexed Targeted iprm-PASEF Assays for MALDI Mass Spectrometry Imaging of Lipids, NAD^+^, and Related Metabolites in CD38 Knockout Mice": Supple Figure 5

**Supplemental Figure 5: Detected fragment ions from the precursor ion isolated at m/z 857.52 and $1/K_0$ 1.4308 ± 0.0093 Vs/cm².** Annotation of the detected ions to known fragment ion m/z from the species PI 36:4(16:0/20:4) confirmed the lipid identity based on characteristic fragments for R-groups Palmitic Acid (m/z 255.23) and Arachidonic Acid (m/z 303.23). Multiple head group fragments, including a prominent ion at m/z 241.01 representing an inositol phosphate ion minus $H_2O$, confirmed the lipid class identity. Pseudo MS/MS spectrum for the isolated species was assembled from detected fragment ions extracted from the iprm mobilogram.

**Pseudo MS/MS for PI 36:4(16:0/20:4)**

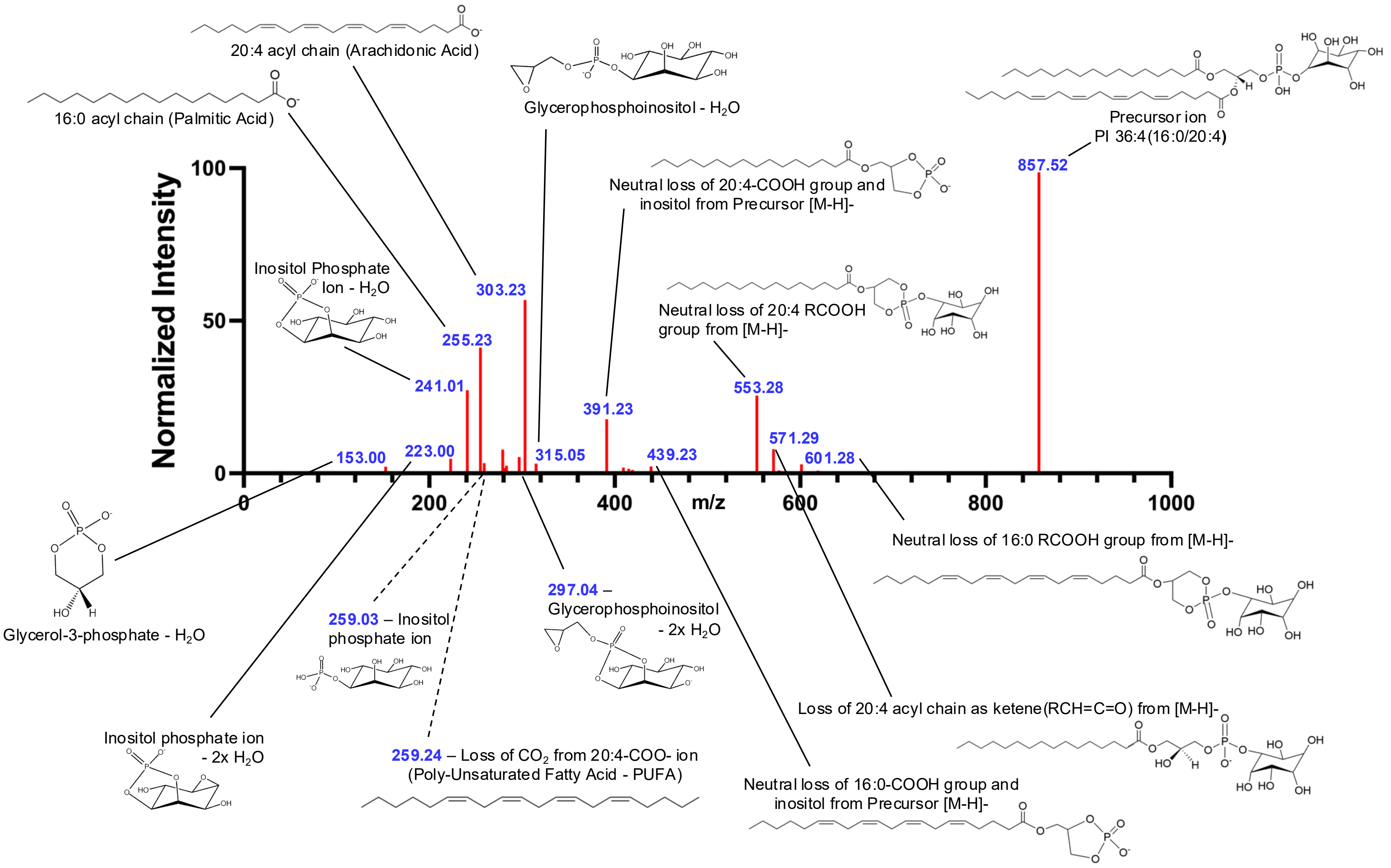
