## Supplementary material for "Multiplexed Targeted iprm-PASEF Assays for MALDI Mass Spectrometry Imaging of Lipids, NAD^+^, and Related Metabolites in CD38 Knockout Mice": Supple Figure 6

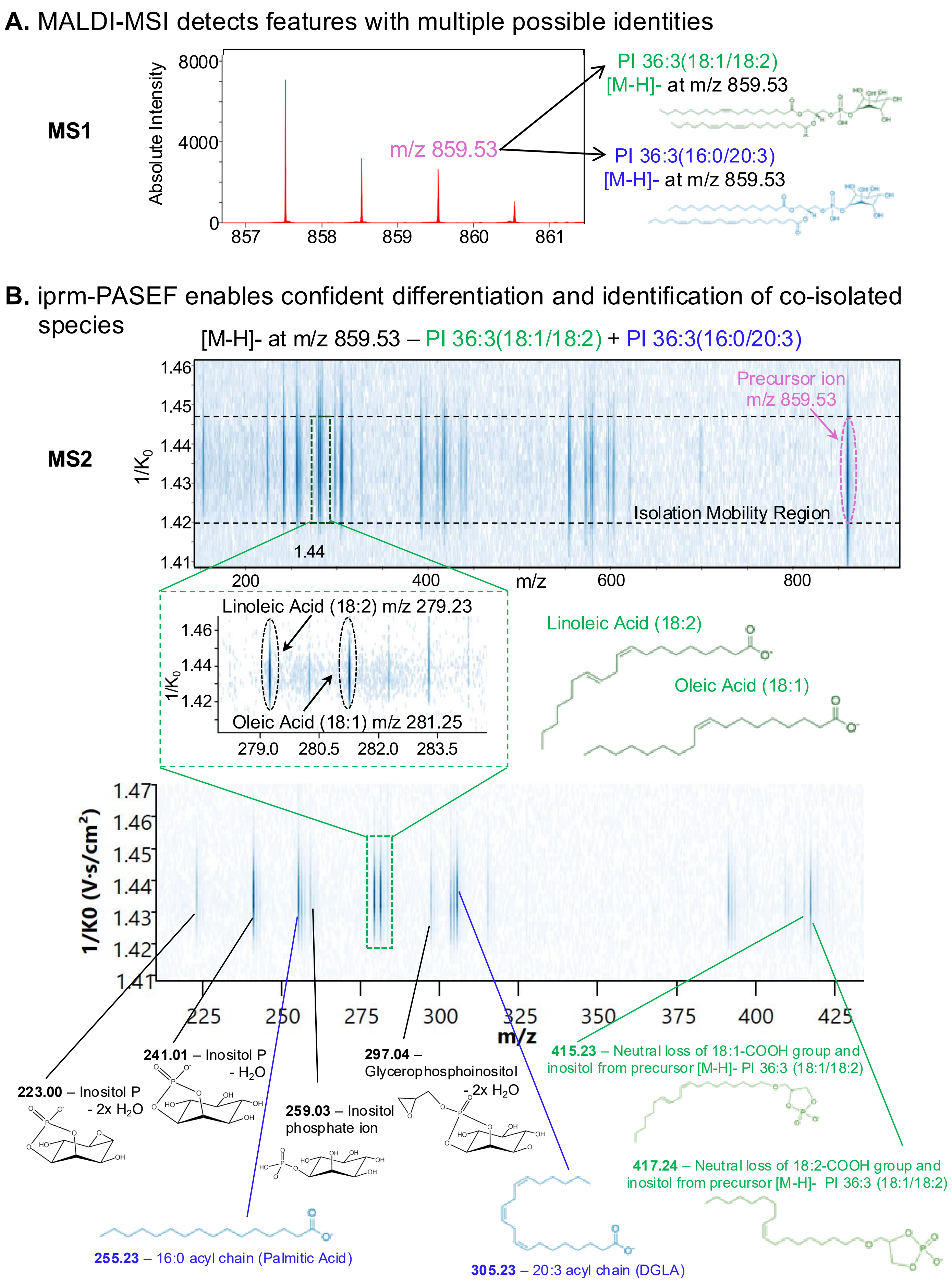

A. MALDI-MSI detects features with multiple possible identities
MS1
Absolute Intensity
8000
4000
0
857
858
859
860
861
m/z 859.53
PI 36:3(18:1/18:2)
[M-H]- at m/z 859.53
PI 36:3(16:0/20:3)
[M-H]- at m/z 859.53
B. iprm-PASEF enables confident differentiation and identification of co-isolated species
[M-H]- at m/z 859.53 – PI 36:3(18:1/18:2) + PI 36:3(16:0/20:3)
MS2
Precursor ion
m/z 859.53
Isolation Mobility Region
1/K0
m/z
Linoleic Acid (18:2) m/z 279.23
Oleic Acid (18:1) m/z 281.25
Linoleic Acid (18:2)
Oleic Acid (18:1)
1/K0 (V·s/cm²)
223.00 – Inositol P - 2x H2O
241.01 – Inositol P - H2O
259.03 – Inositol phosphate ion
297.04 – Glycerophosphoinositol - 2x H2O
415.23 – Neutral loss of 18:1-COOH group and inositol from precursor [M-H]- PI 36:3 (18:1/18:2)
417.24 – Neutral loss of 18:2-COOH group and inositol from precursor [M-H]- PI 36:3 (18:1/18:2)
255.23 – 16:0 acyl chain (Palmitic Acid)
305.23 – 20:3 acyl chain (DGLA)

**Supplemental Figure 6: Isomer resolution of co-fragmented species. A.** iprm-PASEF was used to isolate a precursor ion [M-H]- at m/z 859.53 in the mobility isolation window $1/K_0$ 1.4334 ± 0.0094 $Vs/cm^2$. This species was preliminarily identified as PI 36:3(18:1/18:2) (green) from MS1 survey scans but also matched the precursor ion m/z for PI 36:3(16:0/20:3) (blue). **B.** The fragmentation pattern of the precursor ion [M-H]- at m/z 859.53 with a zoom on the m/z range 279.0-283.5 contained two detected characteristic fragment ions for R-groups Oleic Acid (18:1, m/z 281.25) and Linoleic Acid (18:2, m/z 279.23), while multiple head group fragments, including a prominent ion at m/z 241.01, representing an inositol phosphate ion minus $H_2O$, confirmed the identification of the lipid species PI 36:3(18:1/18:2). Additional fragment ions at m/z 415.23 and 417.24 matched to fragments containing the PI headgroup and tails from the precursor [M-H]- PI 36:3(18:1/18:2). However, further investigation of additional fragment ions across a broader m/z range revealed strong signals for Palmitic Acid (16:0, m/z 255.23) and Dihomo-γ-linolenic Acid (DGLA) (20:3, m/z 305.23). This, along with other fragments containing acyl tails from PI 36:3(16:0/20:3), revealed that the precursor ion [M-H]- PI 36:3(16:0/20:3) was co-isolated with the precursor ion [M-H]- PI 36:3(18:1/18:2) and co-fragmented. Specific and accurate quantification can still be achieved by using characteristic fragment ions for each individual lipid.
