## Supplementary material for "Multiplexed Targeted iprm-PASEF Assays for MALDI Mass Spectrometry Imaging of Lipids, NAD^+^, and Related Metabolites in CD38 Knockout Mice": Supple Figure 7

### Supplemental Figure 7

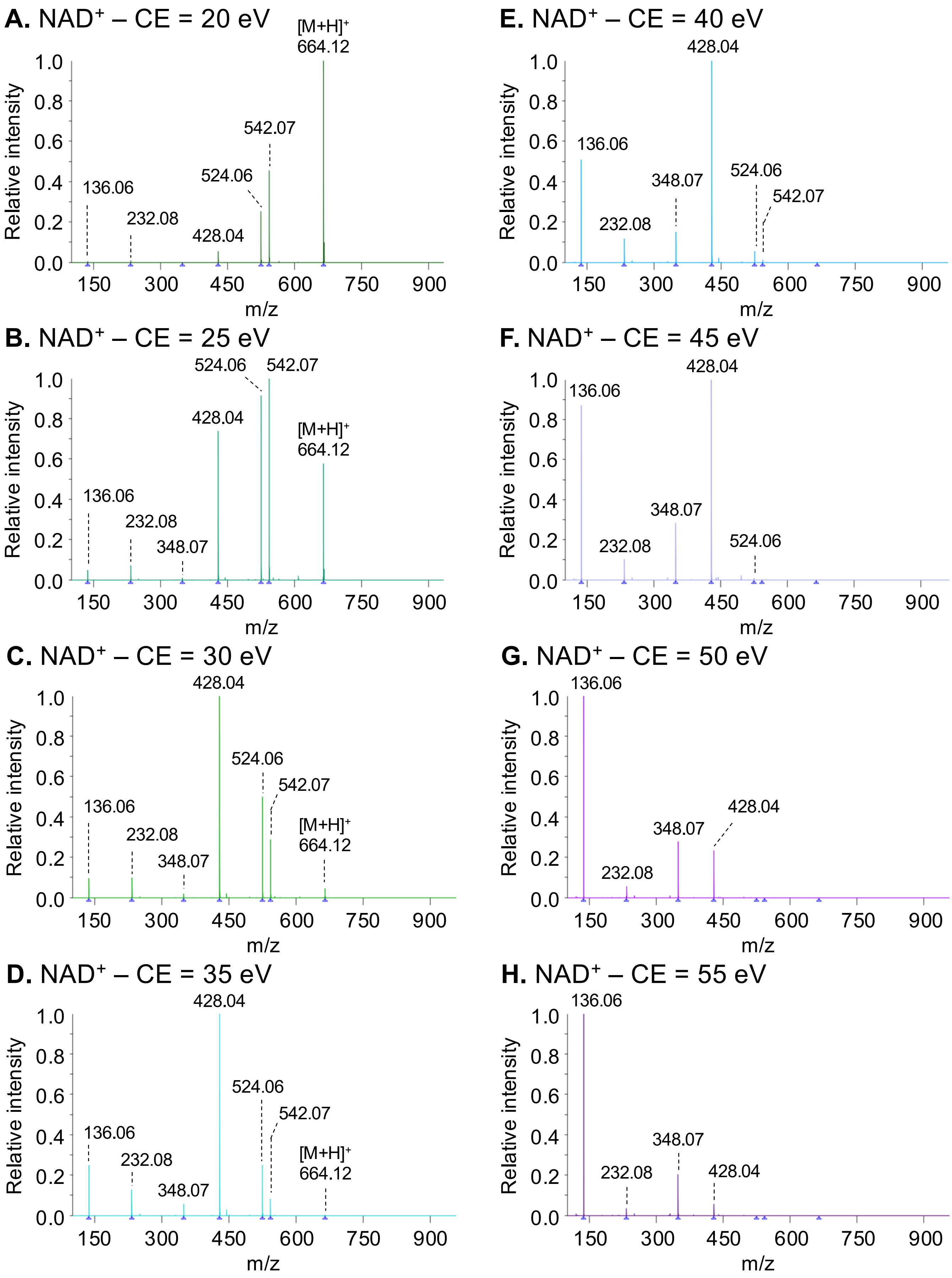

### Supplemental Figure 7

#### I. NAD$^+$ precursor and fragment ion quantification

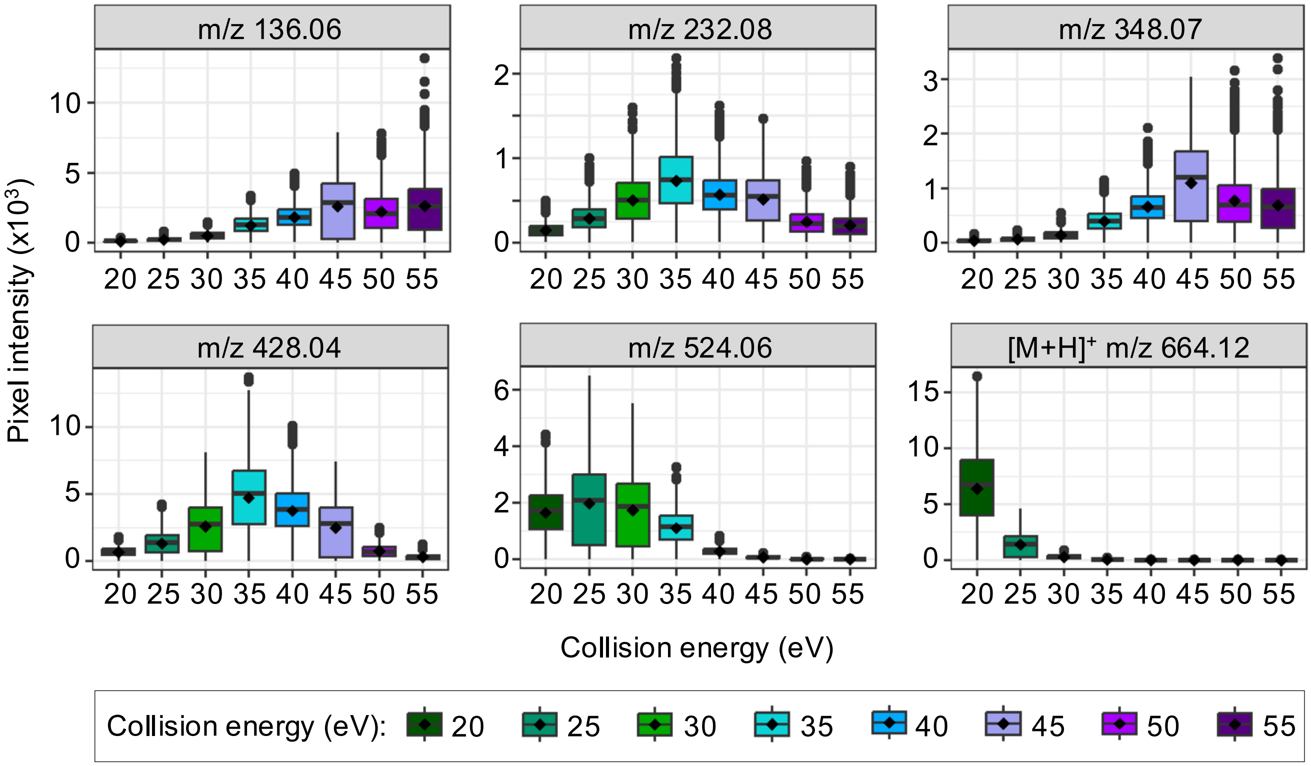


**NAD$^+$ – m/z 664.12**

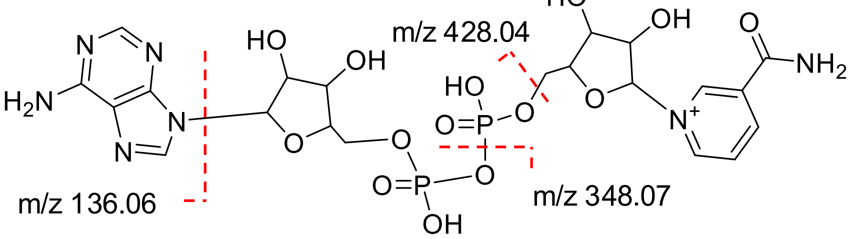


**Supplemental Figure 7: Evaluation of the impact of the collision energy value on NAD$^+$ standard fragmentation pattern.** Synthetic NAD$^+$ standard was spotted on glass slides (200 µM) and DHAP matrix was sprayed for iprm-PASEF analyses in positive mode with precursor ion $[M+H]^+$ at m/z 664.12 and $1/K_0$ 1.12 Vs/cm$^2$. Collision energy (CE) was ramped from 20 eV to 55 eV. A-H. MS/MS spectra of NAD$^+$ standard. I. Boxplots showing the distribution of the pixel intensities of NAD$^+$ fragment ions at m/z 136.06, m/z 232.08, m/z 348.07, m/z 428.04, and m/z 524.06 as well as precursor ion at m/z 664.12 at the different CE tested (see **Supplemental Table 6**). Empty pixels were excluded. The black diamond represents the mean value. Points beyond 1.5 × IQR of the quartiles are shown individually and represent 0–6.3% (median 0.7%) of non-zero pixels per ion and collision energy. Standards were deposited by pipette and dried before matrix was sprayed. This revealed that CE = 20 eV was inefficient to fully fragment the precursor ion: it remained the most abundant ion, while CE ≥ 35 eV was enough to do so. At CE = 25-30 eV, both the precursor ion and the fragment ions were detected. In addition, the fragmentation pattern changed when increasing the CE: m/z 524.06 was the most abundant ion at CE = 25 eV, while it shifted to m/z 428.04 at CE = 30-45 eV and m/z 136.06 for CE ≥ 50 eV.
