## Supplementary material for "Multiplexed Targeted iprm-PASEF Assays for MALDI Mass Spectrometry Imaging of Lipids, NAD^+^, and Related Metabolites in CD38 Knockout Mice": Supple Figure 8

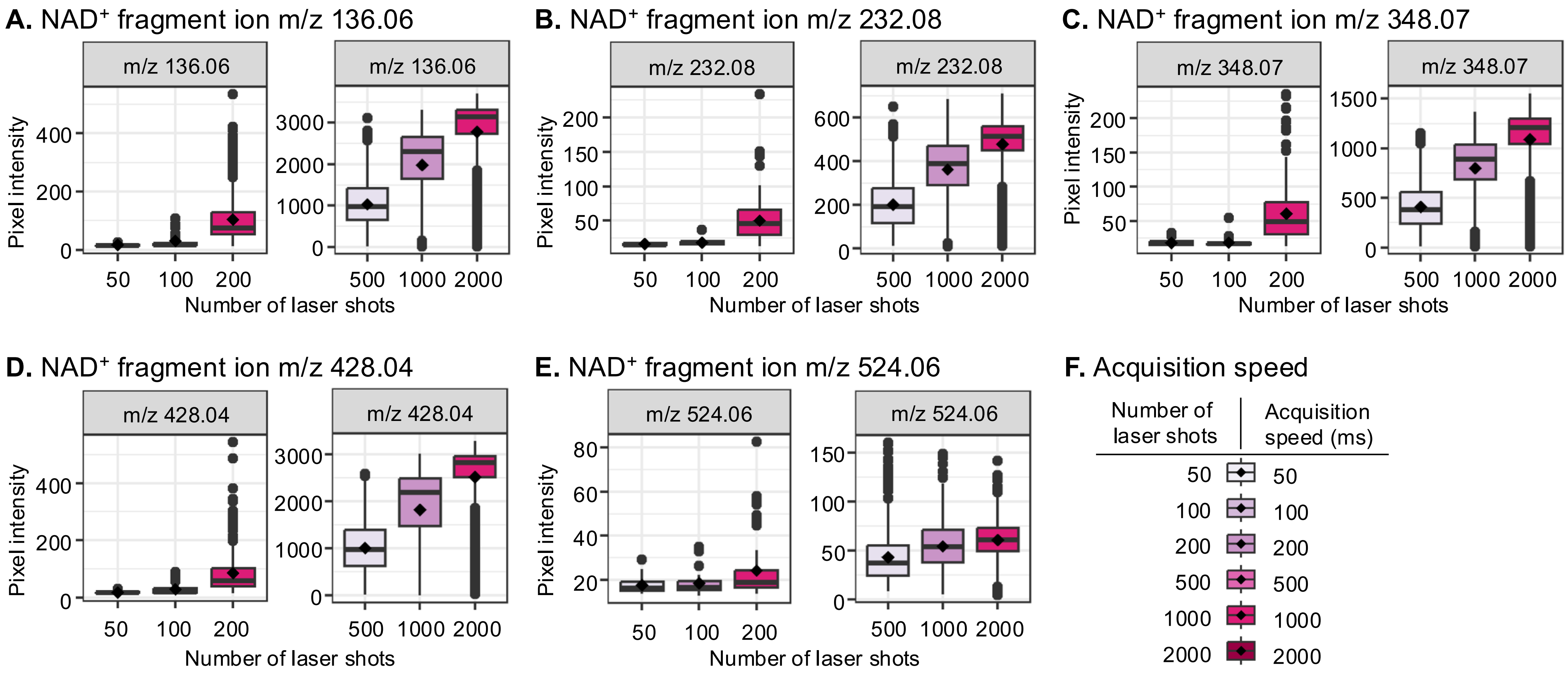


**Supplemental Figure 8: Evaluation of the impact of the number of laser shots on NAD$^+$ standard fragmentation pattern.** Synthetic NAD$^+$ standard was spotted on glass slides (200 µM) and DHAP matrix was sprayed for iprm-PASEF analyses in positive mode with precursor ion [M+H]$^+$ at m/z 664.12 and $1/K_0$ at 1.12 Vs/cm$^2$. The number of laser shots was ramped from 50 to 2000 at 1 kHz. **A-E.** The boxplots show the distribution of the pixel intensities of NAD$^+$ fragment ions at m/z 136.06, m/z 232.08, m/z 348.07, m/z 428.04, and m/z 524.06 for the different laser shot numbers applied (see **Supplemental Table 7**). Empty pixels were excluded. The black diamond represents the mean value. Points beyond 1.5 × IQR of the quartiles are shown individually and represent 0–13.7% (median 4.7%) of non-zero pixels per ion at ≥200 laser shots; at 50 and 100 shots, too few pixels contained signal (9–33 per ion) for outlier percentages to be meaningful. Standards were deposited by pipette and dried before matrix was sprayed. At 50 and 100 laser shots, NAD$^+$ fragments were barely detected. Above 200 shots, the intensity of the fragment ions increased upon enhancing the number of laser shots. **F.** Impact of the number of laser shots on the acquisition speed at 1 kHz laser repetition rate. 200 laser shots per pixel were used for tissue imaging as a compromise between fragment ion intensity and acquisition time.
