## Supplementary material for "Multiplexed Targeted iprm-PASEF Assays for MALDI Mass Spectrometry Imaging of Lipids, NAD^+^, and Related Metabolites in CD38 Knockout Mice": Supple Figure 9

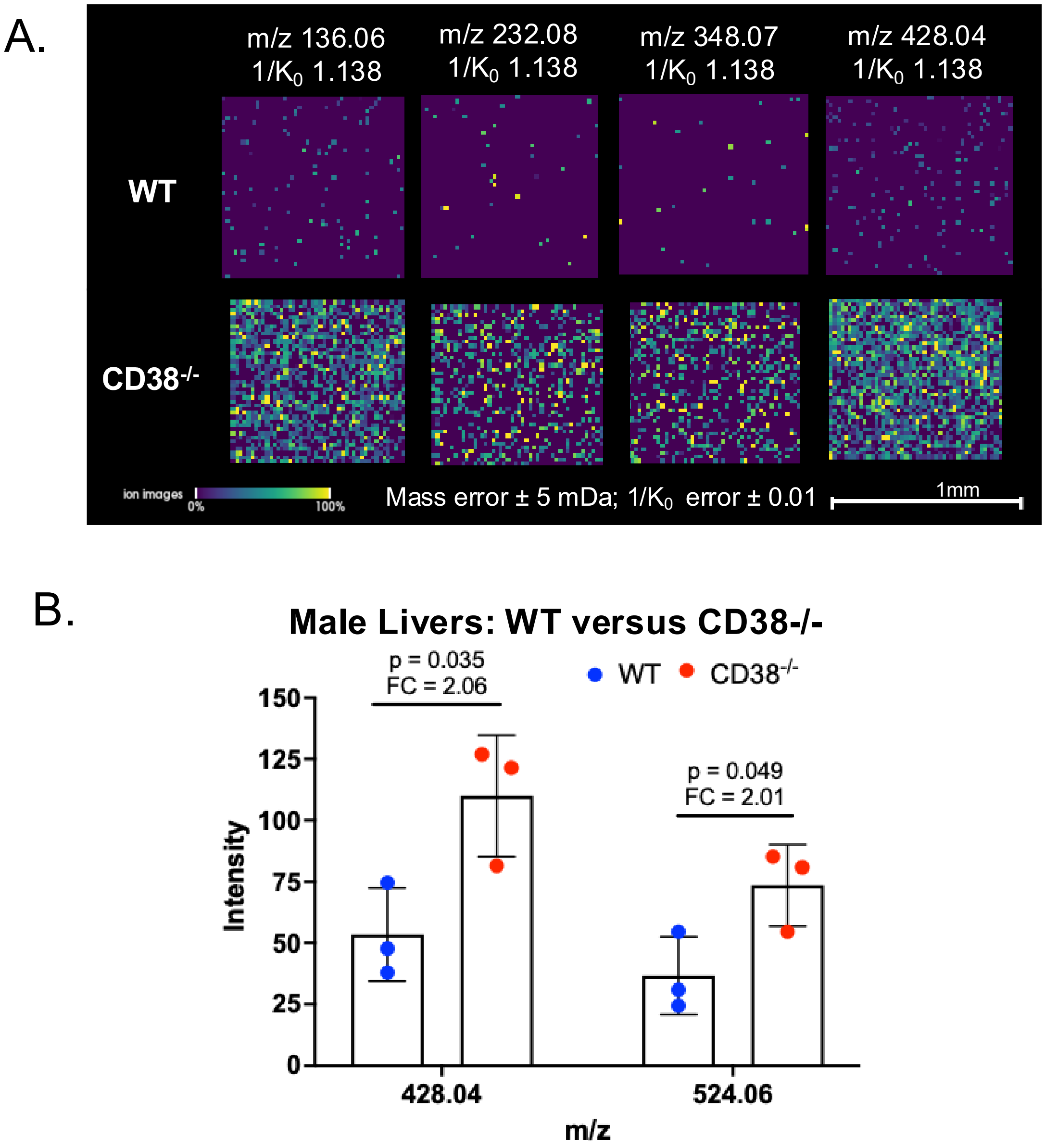


**Supplemental Figure 9**: **Spatial Distribution and Quantification of endogenous NAD$^+$ in livers of male CD38$^{-/-}$ and WT mice via iprm-PASEF.** Spatial distribution of the four NAD$^+$ fragment ions detected by iprm-PASEF in one WT and one male CD38$^{-/-}$ mouse liver replicate show homogeneous distribution of NAD$^+$ when detected in CD38$^{-/-}$ tissue sections and low abundant signal in WT animals. B. Quantification of NAD$^+$ abundance in WT and CD38$^{-/-}$ liver sections in 3 male animals per genotype shows significant increases of fragments m/z 428.04 and m/z 524.06, corresponding to the endogenous NAD$^+$ precursor ion [M+H]$^+$ at m/z 664.12 & $1/K_0$ 1.1224-1.1400 Vs/cm$^2$ in situ during imaging experiments with a fold change of ~2x compared to WT (see **Supplemental Table 4).** For each animal, the non-normalized intensity of each fragment ion was averaged across all non-zero pixels within an ROI selected from the area of highest NAD$^+$ MS1 signal and per-animal means were compared by two-sided Student's t-test. Student's t-test p-value < 0.05 (n = 3 individual mice/biological replicates per condition).
