## Supplementary material for "Multiplexed Targeted iprm-PASEF Assays for MALDI Mass Spectrometry Imaging of Lipids, NAD^+^, and Related Metabolites in CD38 Knockout Mice": Supple Figure 10

### Supplemental Figure 10

**A.** MS1 spectrum of unlabeled $NAD^{+}$ standard

NAD+ – m/z 664.12

Positive mode – DHAP

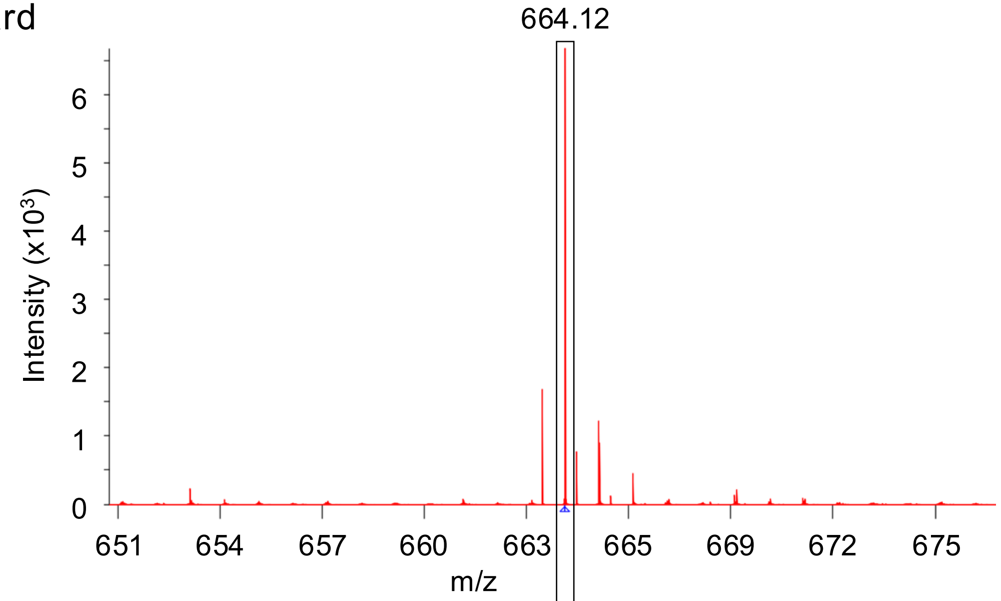


**B.** MS1 spectrum of stable isotope-labeled $NAD^{+}$ standard

$^{13}C_5$-NAD+ – m/z 669.13

Positive mode – DHAP

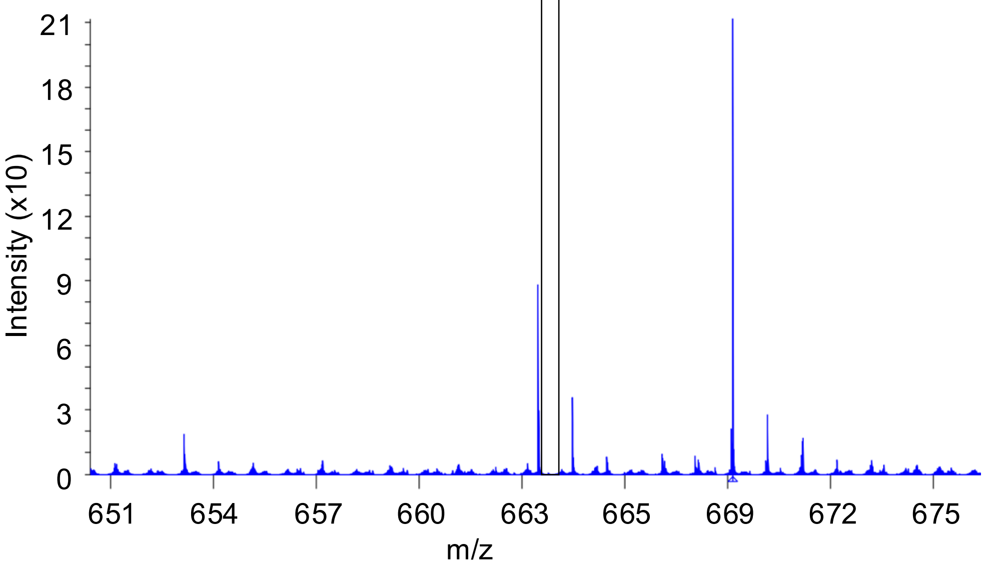

### Supplemental Figure 10

**C.** iprm-PASEF of unlabeled NAD⁺ standard

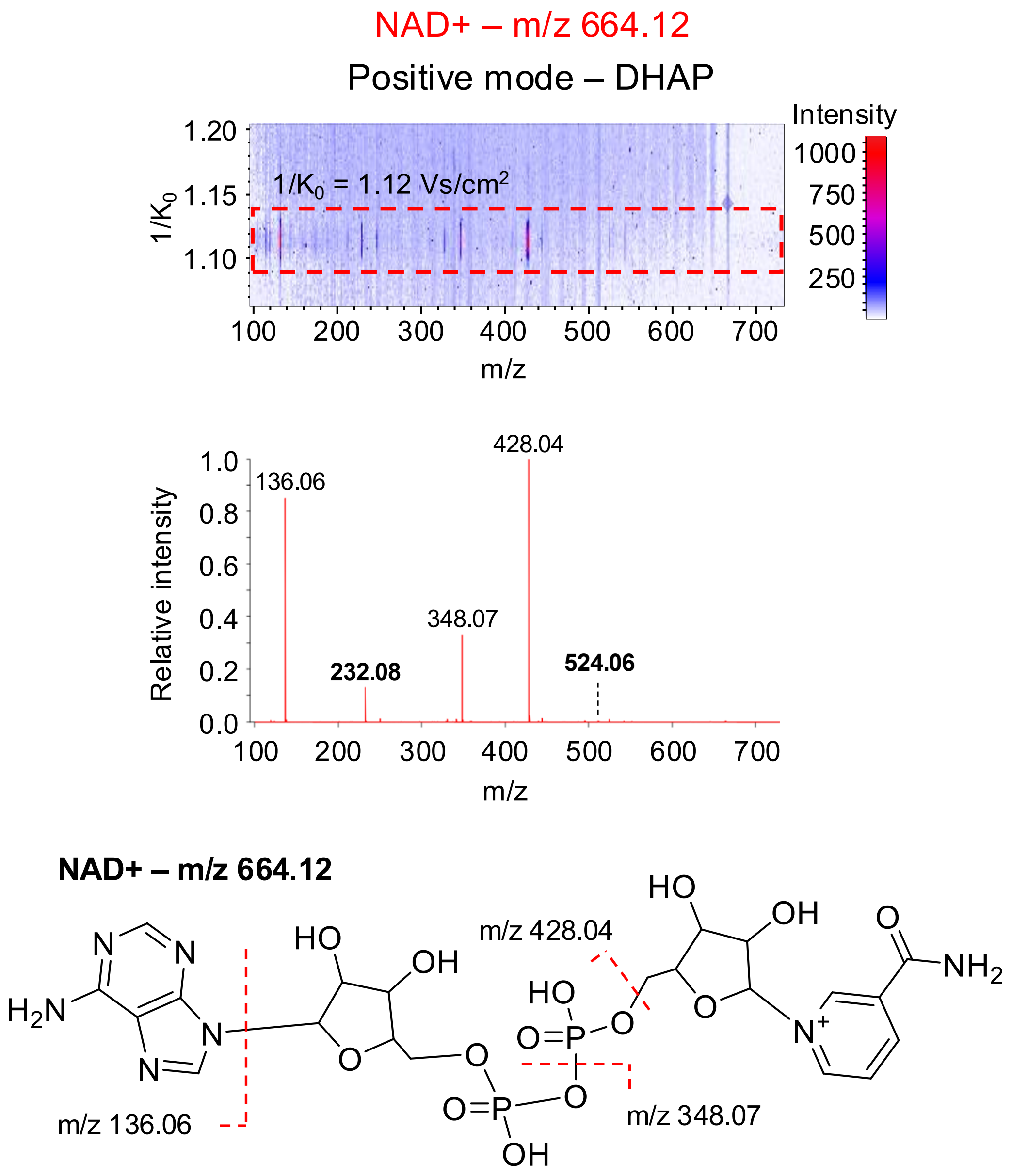


**D.** iprm-PASEF of stable isotope-labeled NAD⁺ standard

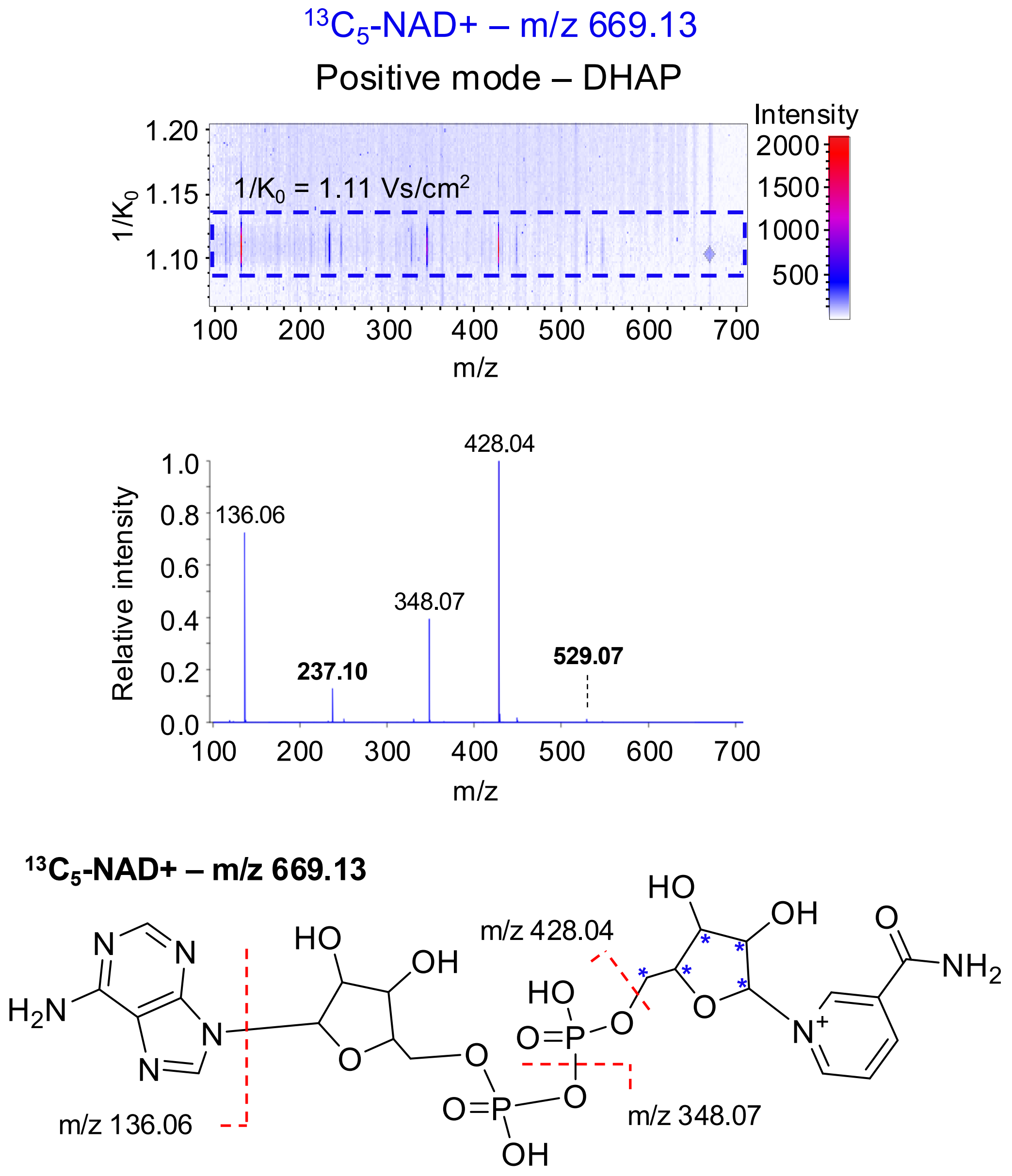

**Supplemental Figure 10: Light and heavy stable isotope-labeled $NAD^+$ standards have the same ion mobility ($1/K_0$) value and fragmentation pattern.** Synthetic light (A,C) and heavy (B,D) stable isotope-labeled $NAD^+$ standards were spotted on glass slides (200 µM) and DHAP matrix was sprayed for survey MS1 and iprm-PASEF analyses in positive mode with light $NAD^+$ precursor ion $[M+H]^+$ at m/z 664.12 and heavy $NAD^+$ precursor ion $[M+H]^+$ at m/z 669.13. A-B. MS1 survey scans are shown for the region m/z 651 – 675 that contains both precursor ions of interest. This shows that the heavy-labeled $^{13}C_5$-NAD+ standard is not contaminated by the light form, confirming that the fragment ions observed by iprm-PASEF analysis specifically results from the fragmentation of the heavy-labelled form. C-D. MS/MS mobility-m/z ratio heatmaps and spectra are displayed. Both standards presented similar $1/K_0$ values of 1.11-1.12 $Vs/cm^2$ as well as similar fragmentation patterns with confident detection of diagnostic ions, *i.e.* m/z 232.08 and m/z 524.06 for the light form as well as m/z 237.10 and m/z 529.07 for the heavy form.
