## Supplementary material for "Multiplexed Targeted iprm-PASEF Assays for MALDI Mass Spectrometry Imaging of Lipids, NAD^+^, and Related Metabolites in CD38 Knockout Mice": Supple Figure 11

### Supplemental Figure 11

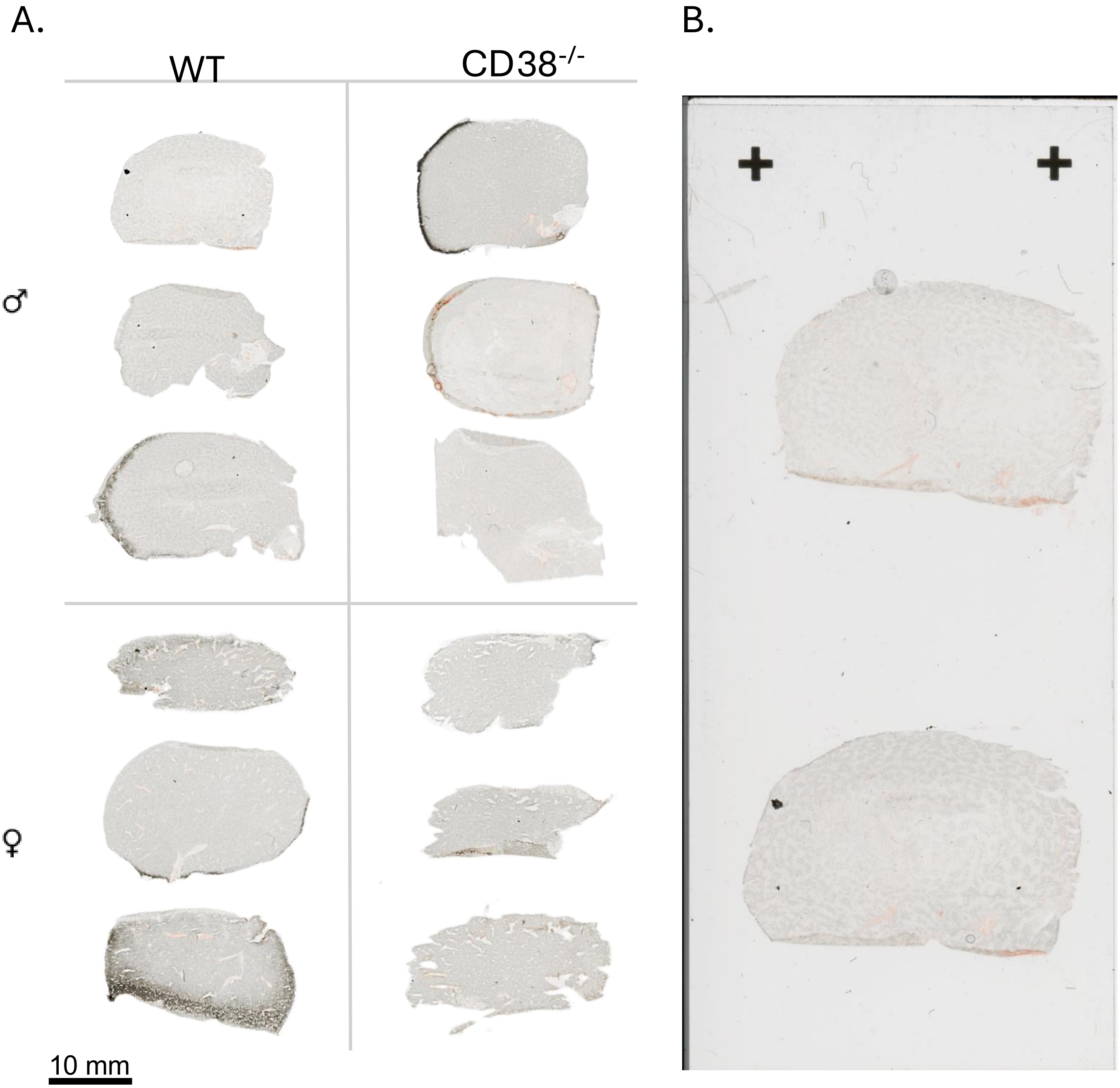


**Supplemental Figure 11: Optical Images of Liver Tissues. A)** Slide scan images of fresh frozen liver tissues used for iPRM-PASEF imaging captured at 2400 DPI for control (WT) and CD38 knock out ($CD38^{-/-}$) mice from male and female. Livers were mounted using carboxymethylcellulose (CMC) and sectioned in the transverse plane through the left lobe attempting to preserve the portal vein structure (lower right) when possible. Scale bar = 10 mm. B) Representative image of serial liver sections mounted on the a standard 25 x 75 mm charged glass microscopy slide.
